# Single-cell transcriptomics reveals evolutionary reconfiguration of embryonic cell fate specification in the sea urchin *Heliocidaris erythrogramma*

**DOI:** 10.1101/2024.04.30.591752

**Authors:** Abdull J. Massri, Alejandro Berrio, Anton Afanassiev, Laura Greenstreet, Krista Pipho, Maria Byrne, Geoffrey Schiebinger, David R. McClay, Gregory A. Wray

## Abstract

Altered regulatory interactions during development likely underlie a large fraction of phenotypic diversity within and between species, yet identifying specific evolutionary changes remains challenging. Analysis of single-cell developmental transcriptomes from multiple species provides a powerful framework for unbiased identification of evolutionary changes in developmental mechanisms. Here, we leverage a “natural experiment” in developmental evolution in sea urchins, where a major life history switch recently evolved in the lineage leading to *Heliocidaris erythrogramma*, precipitating extensive changes in early development. Comparative analyses of scRNA-seq developmental time courses from *H. erythrogramma* and *Lytechinus variegatus* (representing the derived and ancestral states respectively) reveals numerous evolutionary changes in embryonic patterning. The earliest cell fate specification events, and the primary signaling center are co-localized in the ancestral dGRN but remarkably, in *H. erythrogramma* they are spatially and temporally separate. Fate specification and differentiation are delayed in most embryonic cell lineages, although in some cases, these processes are conserved or even accelerated. Comparative analysis of regulator-target gene co-expression is consistent with many specific interactions being preserved but delayed in *H. erythrogramma*, while some otherwise widely conserved interactions have likely been lost. Finally, specific patterning events are directly correlated with evolutionary changes in larval morphology, suggesting that they are directly tied to the life history shift. Together, these findings demonstrate that comparative scRNA-seq developmental time courses can reveal a diverse set of evolutionary changes in embryonic patterning and provide an efficient way to identify likely candidate regulatory interactions for subsequent experimental validation.

## INTRODUCTION

Most metazoan life cycles contain intermediate stages that are ecologically distinct from adults. In many clades, this has resulted in the evolution of contrasting anatomical, physiological, and behavioral traits between stages in the life cycle (Garstang 1928; Thorson 1950; Strathmann 1985; Nielsen 1998; Raff and Byrne 2006; Formery and Lowe 2023). Host-specific stages of parasites, insect larvae, amphibians, and diverse marine invertebrates are often so different from adults that they are unrecognizable from the earlier stages of the same life cycle. In some clades, the evolution of these intermediate stages is remarkably labile, such that closely related species with very similar adult morphology differ profoundly earlier in the life cycle. These cases likely reflect shifts in natural selection that operate on intermediate phases of the life cycle but not on adults. Numerous adaptations related to larval dispersal, feeding, predator avoidance, and abiotic factors have been documented. Yet, it remains largely unknown how developmental mechanisms known to pattern body organization at two distinct stages of the life cycle can become decoupled to allow effective responses to changing selective regimes.

The sea urchin genus *Heliocidaris* provides a valuable system for studying how developmental patterning becomes decoupled across life stages due to a combination of three salient features. First, the genus contains closely related species with highly divergent life histories and a known polarity of change. Second, the selective changes responsible for the life history shift are clear. And third, developmental mechanisms responsible for patterning the ancestral life history are well defined and organized into a developmental gene regulatory network (dGRN). Taken together, these features have made *Heliocidaris* a productive model for understanding genomic and developmental responses to large changes in stage-specific natural selection and their impact on life history evolution (Wang et al. 2019; Davidson et al. 2022a; Davidson et al. 2022b; Devens et al 2023).

*Heliocidaris* illustrates how a shift in selective regimes can rapidly drive extensive changes in intermediate stages (Figure 1A). Representing the ancestral state, *H. tuberculata* produces small (∼100 µm diameter) eggs that develop into complex larvae that feed on phytoplankton for several weeks before achieving sufficient mass to complete metamorphosis. Representing the derived state, *H. erythrogramma* produces much larger eggs (∼430 µm diameter) in greatly reduced numbers, a classic life history trade-off (Stearns 1992). While this ∼100-fold increase in maternal provisioning might seem simple, its impact on other traits has been profound. The larva of *H. erythrogramma* is anatomically highly divergent from *H. tuberculata* (Williams and Anderson 1975; Figure 1D). Unsurprisingly, it has lost the ability to feed: the gut and feeding structures are vestigial, presumably due to relaxed selection. In addition, metamorphosis occurs in just 3.5 days (Williams and Anderson 1975), a reduction of >75% in the duration of the pre-metamorphic phase of the life cycle. This enormous acceleration of early development seems unlikely to be the result of relaxed selection. Instead, the combination of high mortality in the plankton, coupled with greatly reduced fecundity due to the egg size-fecundity trade-off, likely imposes strong directional selection to reduce time to metamorphosis (Wray 2022).

**Figure 1.**
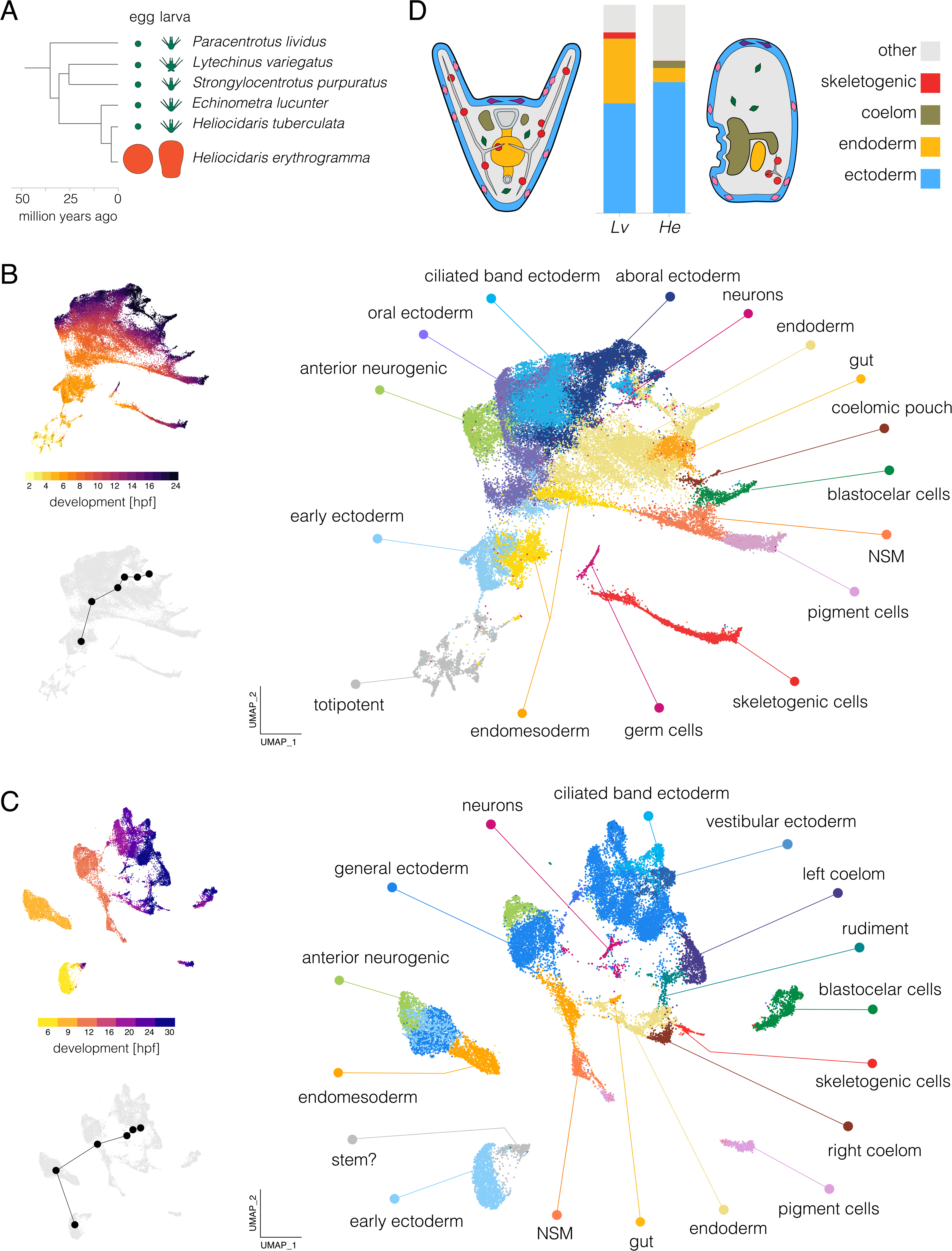
Single-cell developmental transcriptomes. **A.** Time-tree of sea urchin species with high quality reference genomes. Egg and larva sizes approximately to scale; green indicates the ancestral life history with small eggs and feeding larvae (planktotrophy) and orange the derived life history with large eggs and non-feeding larvae (lecithotrophy). Egg and larva sizes from Mortensen 1921, Emlet et al. 1987, Williams and Anderson 1975; topology from Láruson 2016; divergence times from Ziegler et al. 2003 and Láruson 2016. B. UMAPs of scRNA-seq developmental time course for *Lv*. The large plot shows cells color-coded by cluster and labelled according to inferred cell types; the two smaller plots show cells color coded by time point (upper) and the centroids of the six time points common to both species (lower plot). **C.** UMAPS of scRNA-seq developmental time course for *He.* Organization parallels panel B. For individual marker gene expression, see Figure S2. Clusters in panels B and C are colored with the same encoding to facilitate comparison between species (some cell types are present only in one species or the other). **D.** Comparison of cell type proportions and larval morphology. Proportions of four cell types in 24 hpf larvae (see Tables S1 and S2 for cell counts at all stages). Simplified diagrams of larvae are not to scale; colors match bar plot.

In this study, we use single-cell transcriptome analysis (scRNA-seq) to investigate how extensive changes in larval anatomy and a >75% reduction in time to metamorphosis were achieved. We evaluated the presence and relative proportion of larval cell types, the timing of cellular differentiation, trajectories of transcriptional states as a proxy for cell lineages, and the co-expression of transcription factors and targets as indicators of specific regulatory interactions. Our results identify a broad delay in the divergence of transcriptional states during early development; changes in the timing, location, and order of cell fate specification and differentiation; and large shifts in the composition of cell types in the larva. In addition, some ancestral interactions within the dGRN are likely conserved in the derived life history, although most show changes in timing or location, and a few appear to have been lost entirely. Together, these analyses reveal evolutionary changes in embryonic patterning mechanisms and larval biology that were not apparent from morphological comparisons or from bulk RNA-seq analyses.

## RESULTS

### Transcriptional states in *He* accurately reflect the evolution of larval morphology

We began by constructing an atlas of early development in *H. erythrogramma* (hereafter, *He*) for comparison with our previous analysis of *L. variegatus* (hereafter, *Lv*) that spanned early cleavage through early larva (Massri et al. 2021). In order to minimize confounds when comparing between species, our approach to generating data followed the earlier study as closely as possible, including rearing embryos at the same temperature, dissociating cells using only slightly species-optimized protocols, and employing the same generation of library construction and sequencing chemistry (see Methods). We collected seven time points of *He* development from a single cross of outbred adults, from late cleavage (6 hours post-fertilization; hpf) through early larva (30 hpf) (see Methods). We recovered sequences from a total of 23,169 cells after filtering (average ∼3,310 cells/time point). Across samples, we obtained reads from ∼1000 genes/cell and ∼2,000-3,000 UMIs per cell (Figure S1). The number of genes detected per cell drops across the stages sampled, likely reflecting differentiation.

We independently applied clustering and dimensional reduction to the published *Lv* data (Massri et al. 2021) and new *He* data using the same workflow. The resulting UMAPs (Figure 1B,C) are colored by cell cluster (larger plot) and developmental time (upper inset). In both cases, early stages are in the lower left (hot colors) and development proceeds up and right to later stages (dark colors); small gray UMAPs show centroids of stages common to both species. As expected, the spread of points increases during development as cells take on distinct transcriptional states. The distribution of cells is nearly continuous for *Lv*, while that of *He* is more fragmented, likely due to less dense sampling (hourly in *Lv* and every 3 hours in *He*). To identify cell clusters in *He*, we drew on published *in situ* hybridization studies and dGRN genes with conserved expression in specific cell types to annotate clusters with provisional identities (Figure S2; see Massri et al. 2021 for marker genes and supporting literature).

Several cell clusters in the *He* larvae (24 and 30 hpf) could be confidently assigned to a corresponding cluster in *Lv* (24 hpf): pigment cells, blastocoelar (immune) cells, skeletogenic cells, endoderm, coelomic pouch, ciliated band ectoderm, generalized ectoderm, anterior neurogenic ectoderm, and neurons (Figure S2). Each was previously shown to be present in early larvae of *He* based on morphology and marker genes (Mortensen 1921; Williams and Anderson 1975; Parks et al. 1988; Bisgrove and Raff 1989; Wilson et al. 2005a; Love et al. 2008; Koop et al. 2017).

Several differences in the UMAPs reflect the highly derived morphology of the nonfeeding *He* larva relative to the ancestral feeding larvae of most sea urchins, including *Lv* (Mortensen 1921; Williams and Anderson 1975; Wray and Raff 1989). Two clusters present in the 24 hour larva of *Lv* appear to be absent from *He*: primary germ cells and stomodeum (mouth). The absence of germ cells is consistent with the evolutionary loss in *He* of unequal cleavage divisions that found the primary germ cells lineage in the ancestral state (Pehrson and Cohen 1986; Oulhen et al. 2019). The lack of stomodeal cells corresponds to the absence of a larval mouth (Mortensen 1921; Williams and Anderson 1975). Conversely, some cell clusters in *He* larvae are not present *Lv*. These clusters are more challenging to identify: their apparent absence in well studied species with the ancestral life history means that there are no described marker genes. One of these clusters may represent a population of persistent pluripotent cells, based on continued proximity in UMAP space to 6 hpf cells even at 30 hpf (labeled “stem?” in Figure 1C). No corresponding cluster is evident in *Lv* (Figure 1B). Other clusters likely consist of cells that contribute to the adult body (vestibular ectoderm and rudiment, Figure 1C), which develops much earlier in *He* (Williams and Anderson 1975; Wray and Raff 1989; Koop et al. 2017). We also found that ectoderm in *He* does not express markers for the oral and aboral territories present ancestrally, consistent with prior studies based on *in situ* hybridization (Haag and Raff 1998; Love and Raff 2006; Koop et al. 2017). Instead, ectodermal gene expression is organized into a somewhat different set of clusters (Figures 1C and S3) that are not obvious 1:1 homologues of ectodermal territories in *Lv* based.

Substantial differences in the proportions of some cell types are also apparent. Because dissociation protocols can result in biased representation of cell types in scRNA-seq libraries, such findings need to be interpreted with caution. We therefore examined these results in light of prominent morphological differences between the larvae of two species, and highlight three differences that likely reflect true evolutionary changes in cell type proportions in early larvae (Figure 1D; Tables S1 and S2). First, endoderm makes up a much smaller fraction of cells in *He* than *Lv* (6.8 *vs* 31.2%), consistent with its reduced and undifferentiated endoderm (Williams and Anderson 1975; Love et al. 2008). Second, the coelomic pouches contain many more cells in *He* than *Lv* (3.4 *vs* 0.01%). This likely reflects the greatly accelerated development of the imaginal adult rudiment, a large fraction of which is composed of the left coelom (Williams and Anderson 1975; Wray and Raff 1989). Third, a much smaller proportion of skeletogenic cells are present in *He* than *Lv* (0.8 *vs* 2.9%). This is consistent with its greatly reduced larval skeleton (Emlet 1995) and antibody localization of the marker protein Msp130 (Parks et al. 1988). The differences in proportions of the last two cell types are so extreme that they are barely visible in one or the other species in Figure 1D.

### Cell fate specification is broadly delayed in *He*

Examination of the UMAPs at earlier stages of development reveals additional differences (Figure 1B,C). We first identified cell clusters corresponding to two functionally significant territories: the anterior neurogenic domain and the primary signaling center. In the ancestral state, the anterior neurogenic domain is located at the animal pole and develops into the primary sensory organ of the larva (Angerer et al. 2011). The anterior neurogenic domain is clear in *He*, with overlapping expression of dGRN regulators *six3*, *foxQ2*, *nkx3.2*, *zic1*, and *acsc* (Figure S3). The primary signaling center is located at the vegetal pole, and produces ligands that initiate a cascade of signaling events that pattern the animal-vegetal axis (Davidson et al. 1998; McClay 2011). In the ancestral state, the primary signaling center is established ∼3 hpf in the precursors of the skeletogenic cells; they expresses genes encoding ligands, including *wnt8, wnt1*, and *delta* (Sherwood and McClay 1999; Sweet et al. 2002; Wikramanayake et al. 2004; Wei et al. 2012). In *He*, these genes are expressed together within a single cluster (Figure 1C, S2), but beginning much later (6-9 hpf). These cells also express *foxA* and other markers of endoderm (Figure S2). These results suggest that the primary signaling center has become physically separated from specification of skeletogenic cells, a surprising reorganization of pivotal early patterning events in the embryo.

Some other clusters in the *He* embryo could not be confidently assigned to corresponding clusters in *Lv*. The earliest time point sampled (6 hpf) consists of a single cluster lacking any distinctive transcriptional signature (provisionally labeled “early ectoderm” in Figure 1C; the adjacent cluster labeled “stem?” consists of cells from later stages). Remarkably, there is no indication of an early population of either skeletogenic mesenchyme or germ cells in *He*, although these are the first two cell types specified in the ancestral state. Not until 16 hpf in *He* is a population of skeletogenic cells evident, a remarkable delay relative to the ancestral state. While a distinct germ cell cluster is clear in *Lv* (Figure 1B), at no time is a distinct group of cells expressing germ cell markers evident in *He*. *nanos2* and *vasa*, two early regulators of germ cells species with feeding larvae (Juliano et al. 2010; Oulhen et al. 2019), are co-expressed at 9 and 12 hpf in *He* in the presumptive endoderm (Figure S2), but expression disappears at later stages. These observations suggest that some early fate specification events are delayed in *He* relative to *Lv*.

To better understand these differences quantitatively, we systematically analyzed the timing of transcriptional states in the two species. We combined reads without integration prior to dimensional reduction (Figure 2A), which allows us to visualize the relative distribution of cells from each species separately within the same manifold. As expected, reads separated entirely by species on the first dimension (UMAP1), while the second dimension (UMAP2) corresponds to developmental trajectories that run in parallel from bottom to top in both species. These trajectories differ in two informative ways. First, the *Lv* trajectory is initially narrow and spreads into a branch-like structure across UMAP1 as transcriptional states progressively diverge during cellular differentiation (Massri et al. 2021), while the *He* trajectory is also initially narrow but expands only modestly across UMAP1 over time. Given that the transcriptomes of the two species are projected from the same manifold, this contrast suggests overall less divergence in transcriptional states at later stages in *He*. Second, the centroids of embryonic time points in *He* correspond most closely to those of earlier time points in *Lv* along UMAP2 (Figure 2B,C; e.g., 6 hpf in *He* is similar to 2-4 hpf in *Lv*), while later stages occupy similar positions. This suggests that transcriptional states in *He* are initially delayed relative to those in *Lv* but eventually “catch up” during gastrulation. This result is consistent with earlier studies that suggested cell fate specification might be delayed in *He* (Wray and Raff 1989; Wray and Raff 1990; Wang et al. 2020; Davidson et al. 2022a).

**Figure 2.**
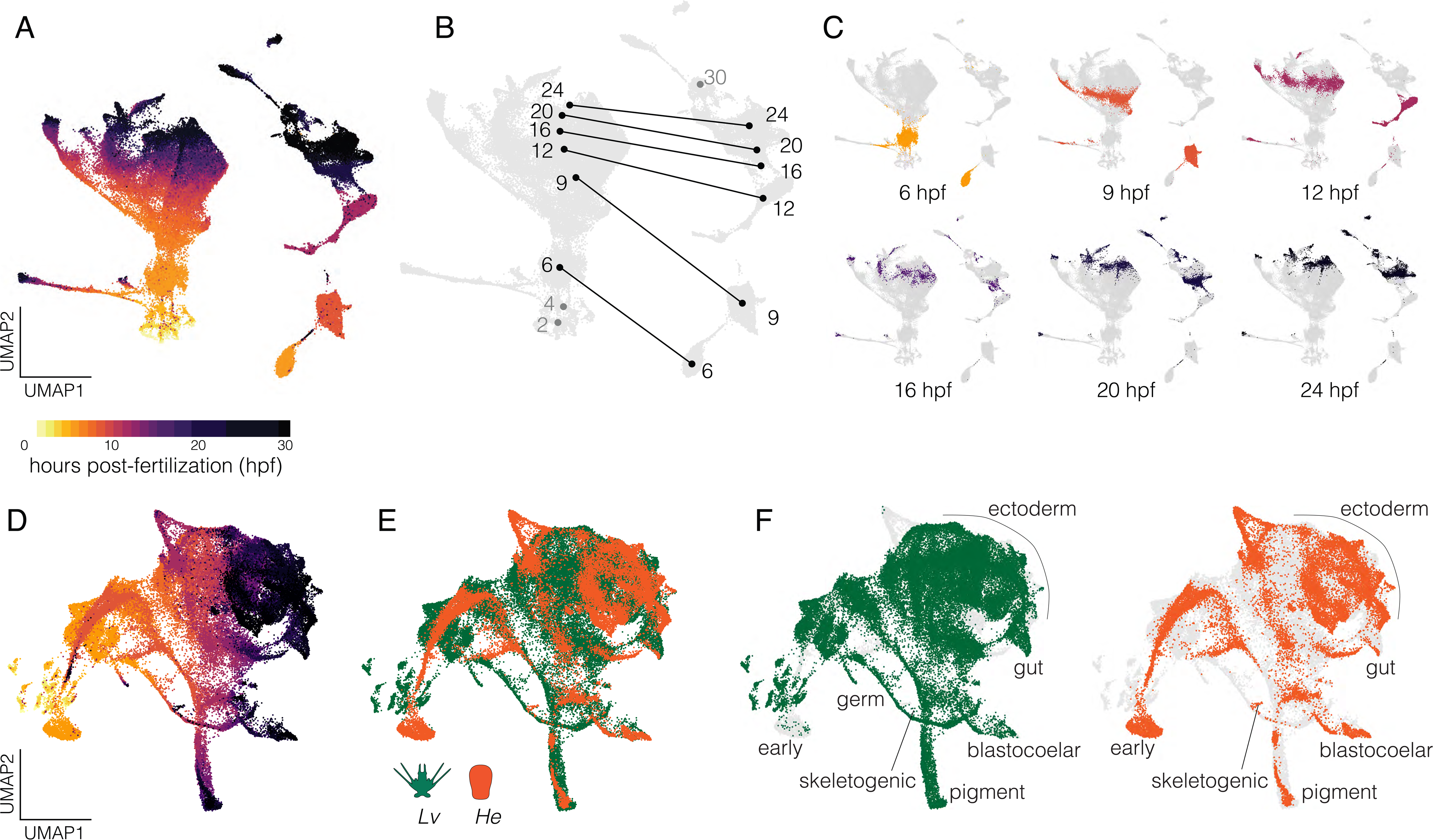
Combined single-cell transcriptomes. **A.** UMAP of combined data from all gene models from both species without integration and cells colored according to developmental stage. The mass on the left is composed exclusively of cells from *Lv* while the more fragmented clumps on the right are composed exclusively of cells from *He*. **B.** Same plot, showing centroids of cells across developmental stages with lines connecting the same time points between species. Numbers correspond to hours post-fertilization (hpf); grey numbers refer to stages sampled in one species only. **C.** Individual UMAPs of shared time points with cells colored according to stage (same encoding as panel A). **D.** UMAP of integrated data from both species incorporating gene models of 1:1 orthologues only. Cells colored by stage (same encoding as panel A). **E.** Same plot with cells colored by species. **F.** Same plot as panel E with each species shown separately and cell identities labeled; light gray cells are from the complementary species.

Next, we integrated reads from 1:1 orthologues of *Lv* and *He* using canonical correlation analysis (Butler et al. 2018) prior to dimensional reduction. The resulting UMAP is presented colored by time (Figure 2D) and by species (Figure 2E). Note that cells from the same time point in the two species generally do not overlap early in development (e.g., “early” in Figure 2F), but that differentiated cells later in development generally do overlap (e.g., “blastocoelar” and “pigment” in Figure 2F). These observations reinforce the inference that diversification of transcriptional states during development take place on different schedules in the two species, with *He* generally lagging at early time points, but later aligning as cells undergo differentiation.

To investigate these evolutionary shifts in timing quantitatively, we used Waddington OT (Schiebinger et al. 2019) to compute transcriptional trajectories for four cell types based on an optimal transport algorithm. We then measured the overall distance between transcriptomes in the two species within each cell lineage (Figure 3A). The most similar time points are indicated by red boxes and plotted in 1:1 aspect ratio in Figure 3B. Points lie predominantly above a line of slope = 1 in Figure 3B in all four cell lineages, indicating that progression through transcriptional states is broadly delayed throughout embryonic development in *He*. We also used a meta-clustering method implemented in CIDER (Hu et al. 2021) to measure overall, rather than lineage-specific, similarity of transcriptomes (Figure S4). Again, results indicate that transcriptional states in *He* lag behind those in *Lv*. This broad transcriptional delay is consistent with a comparison of developmental stages based on morphology, which also shows a delay in *He* (Figure 3C). Because the two species were reared at the same temperature, these rate differences are likely genetically based.

**Figure 3.**
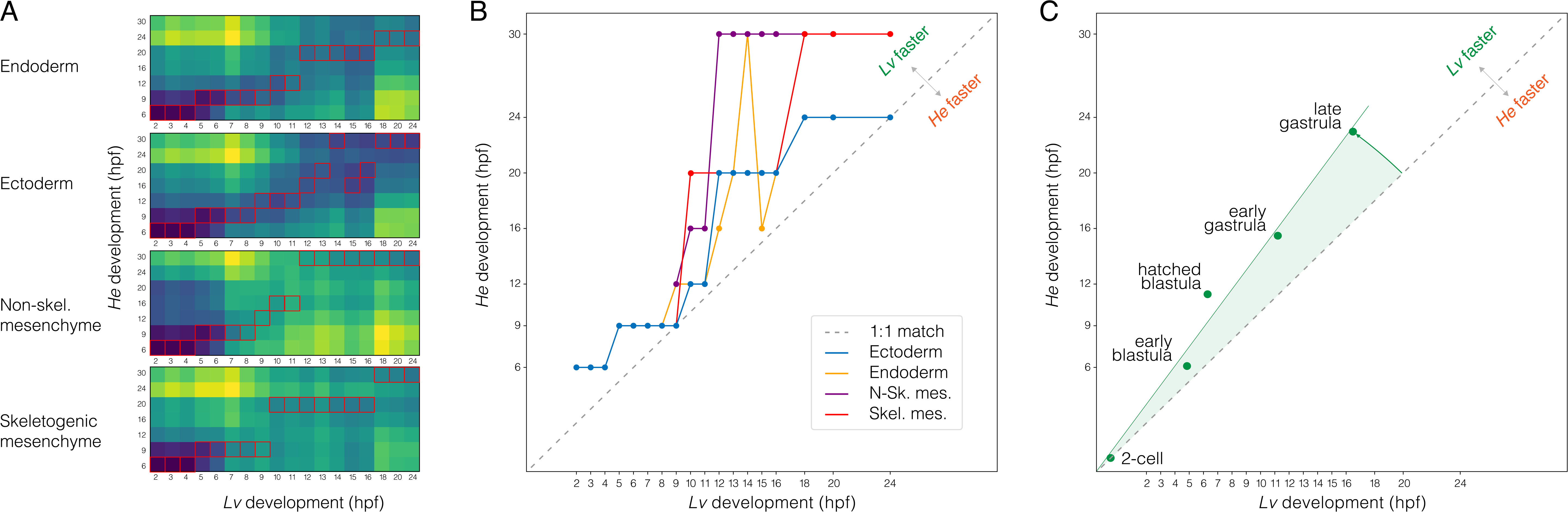
Temporal shifts in transcriptomes. A. Heatmaps showing degree of similarity among scRNA-seq transcriptomes (1:1 orthologues only) for four different embryonic cell lineages. Assignment of cells to lineages is based on optimal transport (see Methods). Red boxes indicate the most similar time point of *He* for each time point of *Lv*. B. Line plots showing developmental time of best matches among transcriptomes in panel A. Note the overall delay in *He*, with most points above the line defined by a slope of 1. C. Line plot showing developmental time of morphogenetic events. Again, there is an overall delay in *He*.

### Differentiation is broadly delayed in *He*

To better understand the timing of expression changes during differentiation, we plotted transcriptional trajectories towards defined differentiated cell states (Figures 4 and S5) following Schiebinger et al. 2019. In Figure 4, each dot corresponds to a cell, with purple and red representing 70% probability of differentiating into a blastocoelar or skeletogenic cell fate, respectively; grey represents transcriptomes predictive of other cell fates and blue indicates uncommitted cells. The triangle is a flattened projection of a high dimensional space and the location of each cell indicates the degree of similarity between its transcriptome and that of two specific differentiated states (top and right apexes) and all other differentiated states (bottom apex). The transcription profile that defines each apex is based on gene expression at 24 hours, when many cell types in *Lv* are approaching a fully differentiated state (Massri et al. 2021). Immediately below each triangle plot is a UMAP showing the location of the same cells.

**Figure 4.**
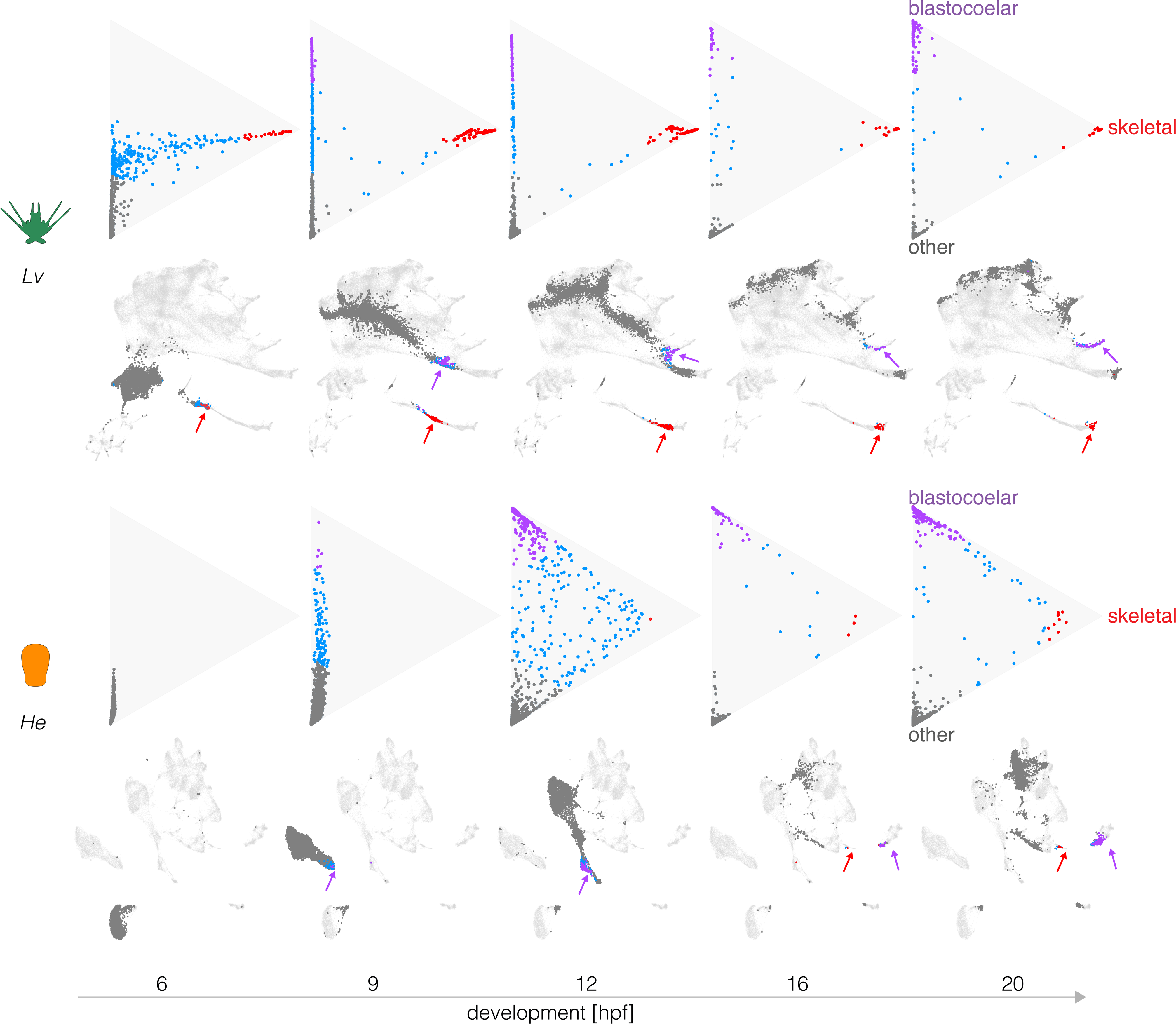
Evolutionary changes in timing of differentiation. Optimal transport was used to predict the likely fate for each cell at five stages, based on transcriptomes at 24 hpf (see Methods). Triangle plots show transcriptomes predictive of blastocoelar cell (green), skeletogetogenic cell (red), or any other cell fate (dark gray); cells with undifferentiated transcriptomes are shown in blue. Corresponding UMAPs are shown below. Note the much earlier differentiation of skeletogenic cells in *Lv* and the slightly earlier differentiation of blastocoelar cells in *He*. See text for additional interpretation.

Figure 4 shows that in *Lv,* many cells take on a transcriptional state predictive of differentiating into a skeletogenic cell as early as 6 hpf (top left triangle plot, red dots). It is not until 9 hpf that a subset of cells are predicted to differentiate into blastocoelar cells (purple dots), consistent with the order of differentiation of these cells in *Lv* (McClay 2011).

Several informative differences are evident in *He*. First, dots are much more tightly bunched at 9 hpf, indicating less divergence in transcriptional states among cells than is the case for *Lv* at the same time and consistent with the delay in diversification of transcriptional states noted earlier (Figures 2 and 3). Second, the order of differentiation is reversed. Blastocoelar cell transcriptomes appear well before those of skeletogenic cells in *He*, whereas skeletogenic cells begin differentiating long before blastoceolar cells in *Lv*. This appears to be due primarily to a shift in skeletogenic cell differentiation, since blastocoelar cells are evident at 9 hpf in both species, while skeletogenic cells appear ∼6 hours later in *He* than *Lv*. Third, at 20 hpf many dots remain far from any apex in *He*, but most dots are at or near an apex in *Lv*. This indicates that more cells remain uncommitted to any specific cell fate in the early larva of *He* than that of *Lv*. Finally, the degree of skeletogenic cell differentiation differs between species. None of the skeletogenic cell transcriptomes reaches the apex in *He, while* many do so in *Lv*, and they begin to arrive much earlier in development (9 hpf). A rather different pattern is seen with blastocoelar cells, where many reach the apex in both species, and this begins earlier in *He* (12 hpf) than *Lv* (20 hpf).

Analysis of other cell types reveals additional evolutionary changes in the timing and degree of differentiation, as well as an example of conservation in timing (Figure S5). Two cell types are noteworthy because changes in their rate of differentiation may be related to the life history shift. Endoderm shows a particularly large delay in the onset of differentiation in *He*: endodermal cells are evident at 9 hpf in *Lv*, but even at 20 hpf none are present in *He* (Figure S5A,B). Coelom shows a less dramatic delay in initial differentiation in *He*, but the number of coelomic cells in *He* overtakes those in *Lv* (Figure S5A). In contrast, blastocoelar and pigment cells show similar overall trajectories in the two species (Figure S5C), indicating that the pace of differentiation within some cell lineages remains relatively unchanged.

Together, these analyses reveal a complex mosaic of evolutionary changes in the timing, order, and degree of differentiation among the two species. While the onset of differentiation is generally delayed in *He*, individual cell types have evolved in distinct ways: blastocoelar cells differentiate earlier in *He* relative to *Lv*, some others are delayed to different degrees (coelom less than skeletogenic cells and gut), and some differentiate at about the same time (pigment cells).

### The order of cell fate specification is altered in *He*

The evolutionary differences in the timing of differentiation noted above are consistent with cell lineage tracing studies in *He* (Wray and Raff 1989; Wray and Raff 1990). However, those studies also suggested that the order of cell fate specification decisions might differ. We therefore reconstructed transcriptional trajectories during development (Chen et al. 2018; Kester and van Oudenaarden 2018; Forrow and Schiebinger 2021) using optimal transport (Schiebinger et al. 2019; Forrow and Schiebinger 2021).

As a positive control, we first evaluated how well transcriptional trajectories based on scRNA-seq data recapitulate actual cell lineages using the published *Lv* time course (Massri et al. 2021), where the cell lineage is well defined by independent methods (McClay 2011). In the resulting directed graph (Figure 5A), nodes correspond to cell clusters and edges connect nodes to their inferred “ancestor” (darker edges indicate higher confidence). This graph contains several features consistent with published analyses of embryonic cell lineages in *Lv* and other sea urchins with feeding larvae (Hörstadius 1973; Pehrson and Cohen 1986; Cameron et al. 1987; Cameron et al. 1990; Ruffins and Ettensohn 1996; Martik and McClay 2017). In particular: skeletogenic and primary germ cells diverge very early; pigment and coelomic cells share a common source population that is distinct from other endomesodermal cells; coeloms, stomodaeum, and gut share a common origin; and neurons derive from both gut and from the anterior neurogenic domain.

**Figure 5.**
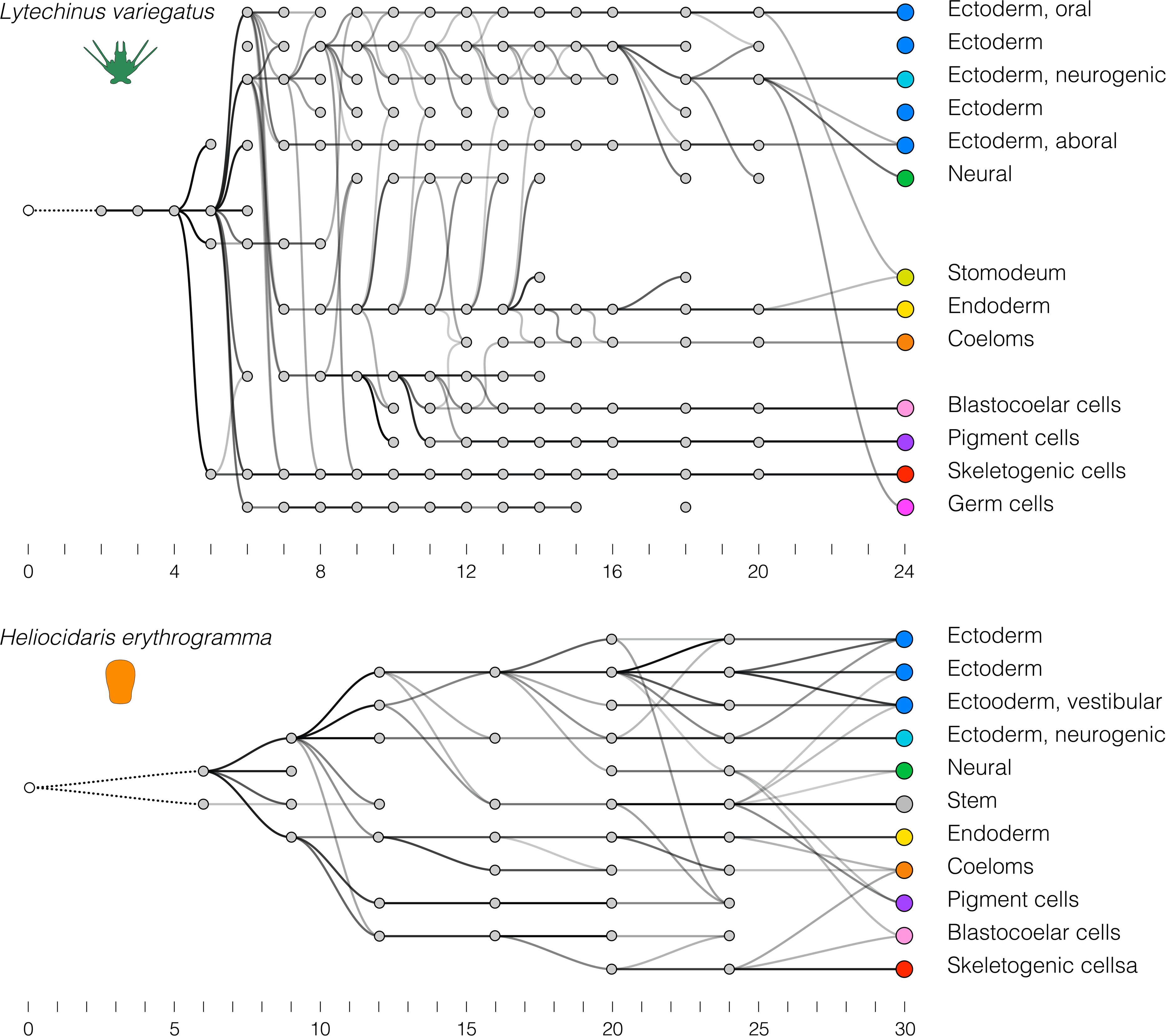
Evolutionary changes in overall transcriptional trajectories. Optimal transport was used to identify “ancestors” for each cell cluster, starting with the final time point (unlike Figure 4, where transcriptomes of individual cells are measured against those of differentiated cells). These trajectories reflect the progressive divergence of transcriptomes among cells during development, and thus are an indirect reflection of cell lineages. **A.** Transcriptional trajectory of *Lv*. **B**. Transcriptional trajectory of *He*.

Some inconsistencies are present, however. Most notably, there is no cluster that corresponds to the four micromeres, the direct ancestors of the skeletogenic and primary germ cell clonal founders. This is likely because cell lineage-specific zygotic transcription is extremely limited at the time the micromeres are present (Ernst et al. 1980) and thus overwhelmed by uniformly distributed maternal transcripts. In addition, the germ cell lineage is discontinuous and shows a late contribution from ectoderm; these are likely artifacts due to their tiny number (8 cells) in proportion to the rest of the embryo at later stages (>1000 cells at 24 hpf). Finally, it should be noted that the ancestor-descendant linkages are in general rather noisy, with several spurious connections. Despite these inconsistencies, the overall topology of the graph resembles the cell lineage as defined by more direct forms of evidence (see citations above).

We then applied the same approach to the *He* scRNA-seq time course (Figure 5B). This graph shares some similarities with that of *Lv*: gut and coeloms are closely related, as are ectodermal territories including the anterior neurogenic domain. Several other features are notably different, however. In *He*, skeletogenic cells are among the last cell clusters to become transcriptionally distinct and are most closely related to blastocoelar cells, while in *Lv* they are one of the first to become transcriptionally distinct and are not related to blastocoelar cells. In addition, pigment cells in *He* are not closely related to blastocoelar cells and become transcriptionally distinct well before they do, while in *Lv* pigment cells and blastocoelar cells derive from a unique common precursor population and simultaneously diverge transcriptionally (Massri et al. 2021). These differences are consistent with the triangle plots (Figures 4 and S5.5). They also imply evolutionary changes in the temporal order and spatial location of fate specification among mesodermal cell lineages (see Discussion).

Other differences in the two graphs are associated with structures or cell types that are present in the larva of one species but not the other. This is apparent in the ectoderm, which is organized anatomically and transcriptionally into somewhat different territories in *He* relative to the ancestral state (Figure S3), with the hugely accelerated appearance of a distinct vestibular ectoderm territory being the most prominent difference (Haag and Raff 1998; Love and Raff 2006; Koop et al. 2017) (Figure 1C). Other notable differences include the apparent absence of endodermally-derived neurons and primary germ cells in *He*.

### scRNA-seq data accurately reflects known regulatory interactions in *Lv*

Results presented above indicate that cells in the embryo of *He* traverse rather different transcriptional trajectories (Figure 4) relative to *Lv*, and that some differentiating cells emerge from distinct precursor populations in the two species (Figure 5). These observations hint at evolutionary changes in underlying regulatory interactions. Two previous studies used scRNA-seq results to infer that specific regulatory interactions present in the sea urchin *S. purpuratus* are absent in the seastar *Patiria miniata* (Foster et al. 2022; Spurrell et al. 2023). We built on this approach, defining criteria for inferring four different evolutionary scenarios: conserved interaction, conserved interaction but with a timing or spatial shift, novel interaction, and loss of interaction (Figure S6).

As positive controls, we first examined experimentally validated regulatory interactions in *Lv*, focusing on the well-studied skeletogenic cell lineage. Figure 6A shows a simplified version of the skeletogenic cell portion of the ancestral dGRN (Kurokawa et al. 1999; Ettensohn et al. 2003; Oliveri et al. 2008; Wahl et al. 2009; Yamazaki et al. 2009; Sharma and Ettensohn 2010; Rafiq et al. 2012; Rafiq et al. 2014). Across the top are the three primary activators of skeletogenic cell-specific transcription, and across the bottom a few of the many effector genes of differentiated skeletogenic cells; between them lie some of the transcription factors that reinforce the differentiated state. We initially focused on *alx1*, which encodes the master regulator of skeletogenic cell specification (Ettensohn et al. 2003; Sharma and Ettensohn 2010), examining interactions involving the two known activators of its transcription (*ets1* and *tgif*) and some of its many known targets (e.g., *dri*, *vegfr*, *sm50*, and *msp130*). We analyzed co-expression of regulators and targets in two ways: as the proportion of cells with co-expression over developmental time (Figure 6B) and by the location of cells with co-expression within the first two dimensions of UMAP space (Figures 7 and S7).

**Figure 6.**
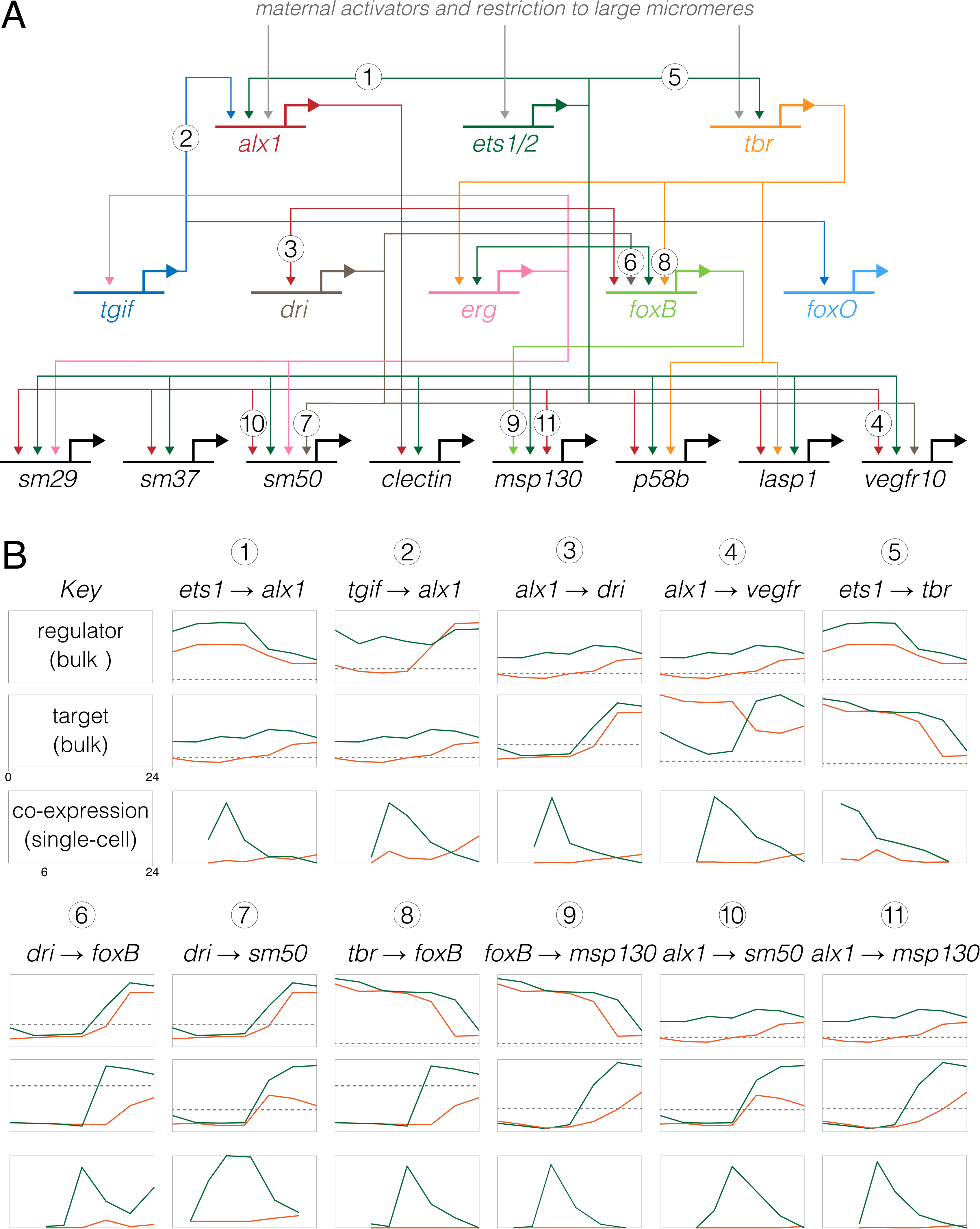
Inference of evolutionary changes in regulatory interactions based on proportion of co-expressing cells. **A**. Simplified version of the skeletogenic portion of the ancestral dGRN present in camarodont sea urchins with feeding larvae (based on Kurokawa et al. 1999; Oliveri et al. 2002, Ettensohn et al. 2003; Oliveri et al. 2008; Rafiq 2012; Rafiq 2014). The three primary activators of skeletogenic-specific transcription (top) feed directly or indirectly into a large set of effector genes, some of which are illustrated (bottom). **B**. Co-expression analysis of 11 experimentally validated regulatory interactions, where *Lv* = green lines and *He* = orange lines. Numbers correspond to interactions in panel A) The top two plots for each interaction show expression of regulator and target based on bulk RNAseq (Israel et al., 2016), with a log2 y-axis and the dashed line indicating very low expression (an average of 5 counts per million reads across time points). The plot directly below shows the proportion of cells that co-express both genes based on scRNA-seq, with a linear y-axis; these time-points begin a 6 hpf, the first time point common to both data sets. Note that y-axes are *not* equivalently scaled because genes have a wide range of expression and co-expression levels. Most gene pairs show a strong peak of co-expression at 9 hpf in *Lv*, which then drops as skeletogenic cells stop dividing while other cell lineages continue to proliferate. In contrast, this peak is notably absent in *He*; instead, co-expression is initially zero or very low at 9 hpf and rises modestly 16-24 hpf.

**Figure 7.**
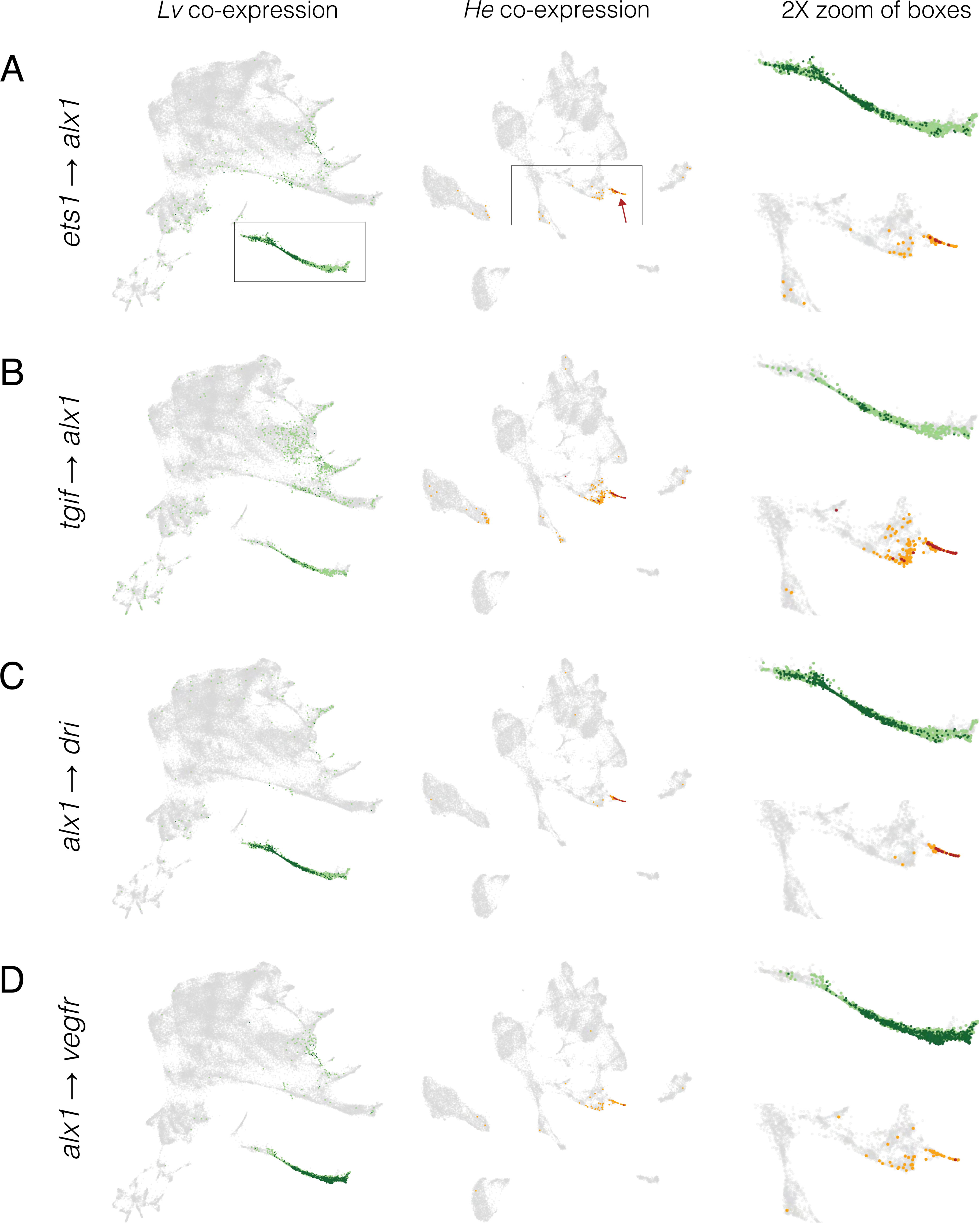
Inference of evolution changes in regulatory interactions based on distribution of co-expressing cells. Co-expression analysis of experimentally validated regulatory interactions. UMAPs show the location of cells with co-expression of indicated regulator and target. *Lv* = green dots and *He* = orange dots; dark colors indicate cells with >2 UMIs for both regulator and target gene; pale colors indicate low co-expressing cells, where one or both genes have 1 or 2 UMIs. Boxes indicate areas shown at 2X in the right-hand column and arrows indicate skeletogenic cells.

Two general points stand out from the *Lv* data (green lines and dots in Figures 6B, 7, S7). First, co-expression of regulator and target occurs at the expected developmental stages. For the six *alx1* interactions shown in Figure 6B, the proportion of cells expressing both regulator and target rises rapidly between 6 and 9 hpf, then declines over time as the skeletogenic cell precursors stop dividing while most other cell lineages continue to proliferate (Martik and McClay 2017). Note that the peaks of co-expressing cells are not evident in the bulk expression of the respective genes (plots immediately above). Many targets of *alx1* show delayed co-expression, with some not yet co-expressed at 6 hpf (e.g., *alx1* → *foxB*) or peaking after 9 hpf (e.g., *alx1* → *sm50*) (Figures 6B, 7, S7). The delay in onset of structural gene expression is consistent with the gap of many hours between skeletogenic cell fate specification and differentiation (Rafiq et al. 2012; Rafiq et al. 2014).

Second, most co-expressing cells are restricted to the skeletogenic cell lineage and the majority of cells in the skeletogenic lineage express both genes (Figures 7, S7). This indicates that co-expression is readily detected despite the sparseness of scRNA-seq data. Both restriction to skeletogenic cells and presence in the vast majority of skeletogenic cells are consistent across many activator → target gene pairs involving *alx1* (Figure 6B, 7, S7) as well as interactions involving other transcription factors within skeletogenic cells (Figure S8). When a transcriptional activator is broadly expressed, co-expression typically involves a specific subset of the cells within its expression domain. For instance, *ets1* and *tbr* are expressed in the endomesoderm as well as in the skeletogenic cell lineage, but co-expression of *ets1-sm32* and *tbr-foxB* are limited to skeletogenic cells (Figure S9).

Overall, these results are consistent with the developmental times and restriction to the skeletogenic cell lineage for these particular regulatory interactions in *Lv*. Figure S10 shows a sampling of co-expression related to regulatory interactions in other embryonic territories. These are also largely consistent with the expected times and locations of experimentally validated regulatory interactions within the ancestral life history. For example, expression of *otx* is quite broad in the embryo, but shows distinct patterns of co-expression with two different targets: in the endoderm, non-skeletogenic mesenchyme, and blastocoelar cells for *gataE*, but just in the endoderm for *endo16* (Figure S10).

### A subset of regulatory interactions may be altered in *He*

Next, we examined the same regulatory interactions in an evolutionary context (Figures 6, 7, 8, S7, S8, and S10; *Lv* = green and *He* = orange). In most cases, co-expression occurs in the same cell lineage in both species. For instance, the ancestral interactions involving *alx1* are reflected as co-expression primarily within skeletogenic cells in *He*. Similarly, *gataE* and *pks1* are co-expressed primarily in the pigment cells (Figure S10). These results are consistent with conservation of ancestral regulatory interactions in *He*. Several differences between species, however, point to evolutionary changes in specific regulatory interactions, including: changes in timing (earlier or later), location (extent or cell lineage), and presence/absence (Figure S6).

The most common differences are in timing of co-expression. The prominent early peak of *alx1* interactions in *Lv* is reduced or entirely absent in *He* (Figure 6B); instead, the proportion of co-expressing cells rises later in *He*, reflecting much later differentiation. This is largely due to a late rise in *alx1* expression (Figure 6B; dashed line indicates < 5 transcripts per million across 3 replicates). Among targets of *alx1* expression, *dri* shows a similar expression profile among species, while *vegfr* shows highly divergent expression; nonetheless, the co-expression time courses and UMAPs are very similar for both interactions. The simplest explanation for these results is that some regulatory interactions take place in *He* but that they are considerably delayed relative to *Lv*. Earlier observations indicating a delay in both specification and differentiation of skeletogenic cells in *He* (Figures 1, 3, 4, 5) are consistent with this interpretation.

Less commonly, an interaction appears to be absent in *He*. For the ancestral interaction *alx1* → *foxB*, no cells at any time contain reads from both genes in *He* (Figure 8). Since both genes are robustly expressed at other times and locations in the *He* embryo, the complete absence of co-expression is not simply a technical issue with detection. Another example is the ancestral interaction *alx1* → *foxO*, where in *He* only one cell across all time points expresses both genes and it contains low UMI counts from each gene (indicated by light orange) (Figure S7). Given that co-expression often occurs in a small number of scattered cells outside the region where a specific interaction is thought to occur (Figures 7, S7, S8, S10), this low level of co-expression of *alx1* and *foxO* in *He* is likely not functionally significant. Other examples involves *tbr*, which is not expressed in skeletogenic cells in *He*, despite being expressed elsewhere in the embryo (Figure S8). The simplest interpretation is that these regulatory interactions do not take place in *He*.

**Figure 8.**
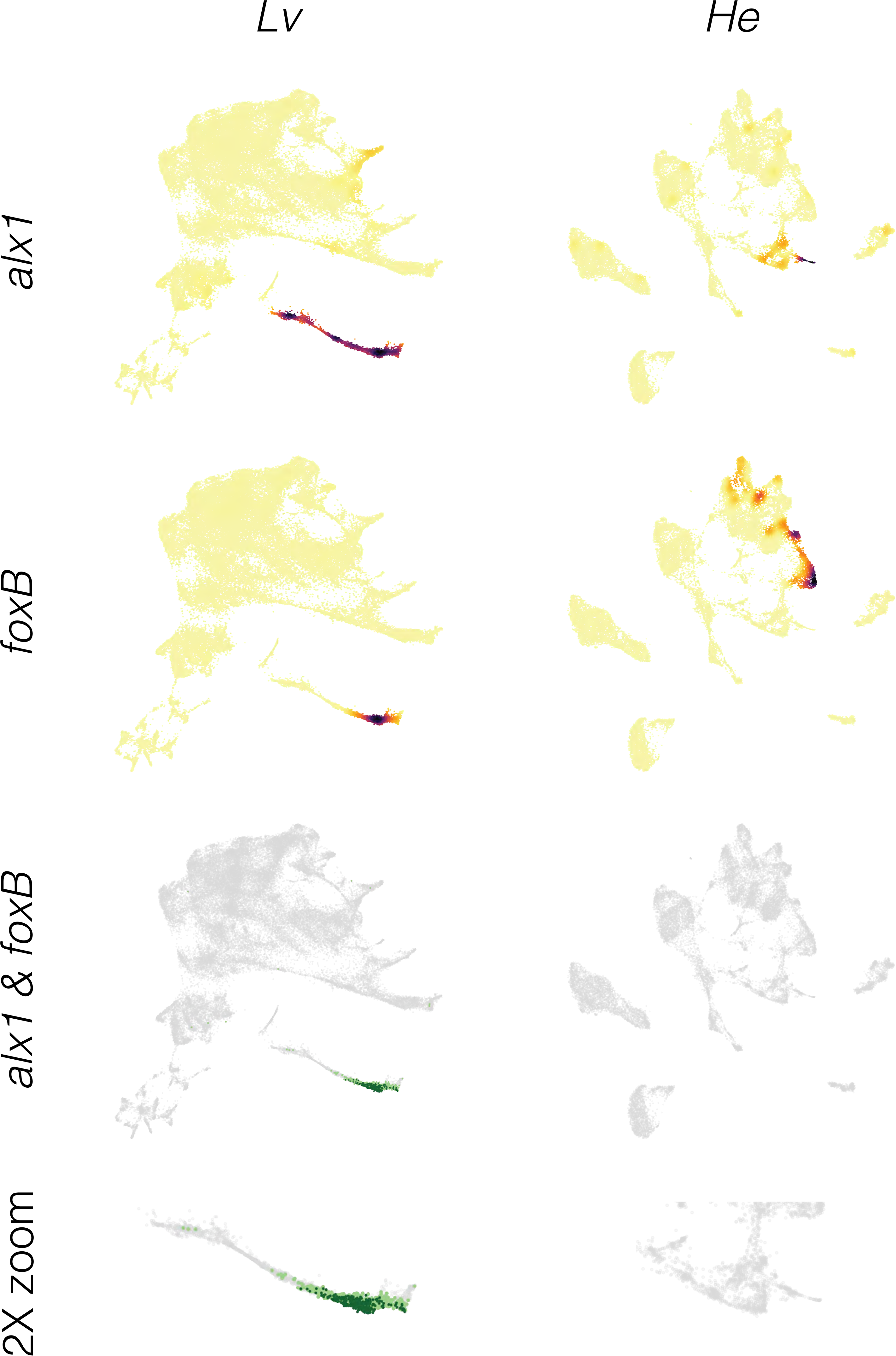
Inference of evolutionary loss of a regulatory interaction. **A.** Density plots showing expression of regulator (*alx1*) and target (*foxB*) genes in both species. Note that both *alx1* and *foxB* transcripts are readily detected in both species. **B.** Co-expression plots. The complete absence of co-expression in *He* suggests that the ancestral *alx1* → *foxB* regulatory interaction has been lost in this species.

Although the focus here has been on skeletogenic cells, the same general findings are evident in other territories (Figure S10). Again, most co-expression of regulator and target in *Lv* corresponds to expected times and locations. When comparing species, most co-expression occurs in the same embryonic territory or cell lineage, but with notable exceptions that suggest specific kinds of evolutionary change (Figure S6). Among these, differences in timing or location are the most common. For instance, co-expression of *otx* and *gataE* occurs throughout the endomesoderm of both species, but is largely confined to 9-12 hpf in *He* while present from 6 to 24 hpf in *Lv*. Similarly, co-expression of *bra* and *foxA* is largely endodermal in both species, but in addition is more extensive in the ectoderm of *He* than in *Lv*. A few additional likely losses of regulatory interactions are also evident. These include *bra* → *apobec* within the endoderm, which may be absent in *He* (Figure S10). Due to the large number of documented interactions within the ancestral dGRN, a comprehensive co-expression analysis throughout the dGRN is beyond the scope of the present study.

## DISCUSSION

Comparisons of single-cell transcriptomes between species have been used to document the presence or absence of cell types (e.g., Levy et al. 2021; Tarashansky et al. 2021; Wang et al. 2021; Alvarez-Campos et al. 2023; Mah and Dunn 2023), but less commonly to understand how developmental mechanisms evolve and contribute to organismal traits. This study used scRNA-seq to examine the evolution of cell fate specification and differentiation in *Heliocidaris erythrogramma* (*He*), a sea urchin with a recently modified life history (Figure 1A) (Raff 1992; Ziegler et al. 2003; Wray 2022). The goal was to gain insights into the developmental basis for massively modified larval morphology and hugely abbreviated pre-metamorphic development. We generated a developmental time course of scRNA-seq data from *He* and carried out comparative analyses with our published data for *Lytechinus variegatus* (*Lv*) (Massri et al. 2021), representing the ancestral life history in sea urchins (McEdward and Miner 2001; Raff and Byrne 2006). This discussion is organized around three broad themes revealed by comparative analyses of the scRNA-seq time courses.

### Evolution of embryonic patterning

The earliest indication that embryonic patterning might be modified in *He* came from observations of cleavage divisions, which differ from the stereotypical pattern in the ancestral life history. In sea urchins with feeding larvae, unequal vegetal cleavage divisions establish the clonal founders of two distinct cell lineages: the germ line (Pehrson and Cohen 1986; Oulhen et al. 2019) and the skeletogenic cells, which also become the primary signaling center of the embryo (Hörstadius 1973; Sherwood and McClay 1999; Sweet et al. 2002; Wikramanayake et al. 2004; Wei et al. 2012). These processes appear to be conserved in *H. tuberculata*, a close relative of *He* (Figure 1A) (Wray and Raff 1988; Love et al. 2007; Morris et al. 2019). Over the next few hours, a series of inductive interactions initiated by the primary signaling center specify other embryonic cell lineages (reviewed in McClay 2011). These critical early patterning events are broadly among sea urchin species with the ancestral life history (McClay 2011; Thompson et al. 2015; Minokawa et al. 2017; Yamazaki et al. 2021) (Figure 1A). In contrast, *He* lacks any early unequal cleavage divisions (Williams and Anderson 1975). Dye-tracer studies reveal a general delay in specification and, specifically, that no early blastomeres are clonal founders of either the germ line or skeletogenic cells (Wray and Raff 1989; Wray and Raff 1990).

The scRNA-seq results reported here confirm this delay and add new information. At 6 hpf, the embryo of *Lv* contains four transcriptionally distinct populations of cells: germ cell precursors, skeletogenic cell precursors, early ectoderm, and endomesoderm (Massri et al. 2021) (Figure 1B). In contrast, the 6 hpf embryo of *He* contains a single population of cells producing transcripts characteristic of undifferentiated epithelium (Figures 1C and S2).

Closer examination of each of the three early patterning events reveals striking changes in early patterning. (1) *Skeletogenic cells*. Both species express *alx1*, which encodes the master regulator of skeletogenic cell fate (Ettensohn et al. 2003), but with a large delay from ∼3 hpf in *Lv* to later than 12 hpf in *He* (Figures 1C, S2 and S7). A distinct skeletogenic transcriptional state is apparent by ∼6 hpf in *Lv but* not until ∼16 hpf in *He*. Even at 30 hpf, skeletogenic cells of *He* are not as differentiated as they are at 24 hpf in *Lv* (Figure 4). (2) *Germ cells*. At no point up to 30 hpf in *He* is there a distinct cell population expressing germ cell markers. Species with the ancestral life history express *nanos2* broadly, but transcripts and protein accumulate exclusively within the small micromeres (Oulhen et al. 2019). Expression of *nanos2* also occurs in *He*, but does not become localized, remaining widespread in endomesoderm up to 30 hpf (Figure S2). The same is true of *vasa*, another germ line marker (Figure S2). (3) *Primary signaling center*. In *Lv*, the skeletogenic precursors between 3 and 6 hpf express genes that encode signaling ligands, including *wnt1*, *wnt8*, and *delta* (Massri et al. 2021). In *He*, however, these genes are co-expressed with markers of endomesoderm (*foxA*, *ism*, *blimp1*; Figure S2) and never with skeletogenic markers, suggesting that the primary signaling center is spatially separated from skeletogenic cell fate specification. In addition, the timing of expression differs in *He*: only *wnt8* is expressed at 6 hpf, while all three transcripts show peak expression at 9 hpf (Figure S2). These observations indicate that the three earliest embryonic patterning events in the ancestral state are all delayed in *He*, and that they have become spatially and temporally separated from each other. This delay is reflected more broadly in the embryo, with transcriptional states in multiple territories diverging later on average (Figures 3, 4, and 5).

The simplest model to explain these observations is that embryonic patterning mechanisms are conserved in *He* but activated later in development. Three lines of evidence suggest that the situation may actually be more complicated. First, the three earliest patterning events are nearly simultaneous in the ancestral state but occur at widely separated times in *He*: the primary signaling center is established prior to 9 hpf, skeletogenic cell fate specification takes place between 12 and 16 hpf, and germ cell fate specification occurs some time after 30 hpf. Second, in a previous study, we showed through perturbation experiments that the earliest regulatory interactions responsible for skeletogenic cell fate specification have been lost in *He* (Davidson et al. 2022a). Germ cell fate specification has not been experimentally investigated in *He*, but *foxY*, which encodes a key regulator of *nanos2* transcription in the ancestral life history (Oulhen et al. 2019), is not tightly co-expressed with it. Third, some populations of larval cell types in *He* derive from different founder cells than in the ancestral condition (Figure 5). In particular, pigment cells and blastocoelar (immune) cells derive from a uniquely shared population of non-skeletogenic mesenchyme cells in *Lv* (Figure 1B) and other species with the ancestral life history (McClay 2011); in contrast in *He* these two cell types derive from spatially and temporally distinct source populations, and instead it is blastocoelar and skeletogenic cells that share a common origin (Figures 1C and 5). Thus, embryonic patterning and cell fate specification appear to be rearranged in a manner inconsistent with a simple conservation-with-delay model.

Importantly, not all embryonic patterning events are delayed in *He*. A striking counter-example is the breaking of left-right symmetry, which occurs before first cleavage (Henry and Raff 1990; Henry et al. 1991). In contrast, the first indication of left-right asymmetry in the ancestral developmental mode occurs in the late gastrula (Duboc et al. 2005; Bessodes et al. 2012). Another accelerated patterning event in *He* involves the early establishment of the imaginal adult rudiment, which begins at about 30 hpf in *He* (Williams and Anderson 1975; Wray and Raff 1989; Koop et al. 2017) but not until several days post-fertilization in the ancestral condition (Lowe et al. 2002; Formery et al. 2022).

In sum, patterning mechanisms in the early embryo of *He* appear to represent a complex mosaic of changes. Three critical early patterning events that are tightly associated with a set of unequal cleavages in the very early embryo of the ancestral state are delayed in *He*, and in addition are dissociated from each other in time and location. In contrast, some other pivotal patterning events are accelerated in *He*. Furthermore, the origins of some larval cell types have been rearranged, likely reflecting changes in embryonic cell lineages. Together, these changes suggest that several modifications have evolved in interactions within the dGRN, as discussed next.

### Evolution of regulatory interactions during development

The ability to assay transcription from single cells provides exciting opportunities to investigate the evolution of transcriptional regulation. In particular, the interaction between a transcriptional activator and a regulatory target should be reflected by co-expression within the same cell. It is important to emphasize that co-expression does not by itself constitute direct evidence: it can reveal a pattern consistent with a regulatory interaction, but experimental evidence is needed to confirm. For this reason, we restrict attention here to gene pairs representing experimentally documented regulatory interactions in the ancestral state, rather than attempting to identify previously unknown interactions. Our current understanding of ancestral dGRN interactions in sea urchins comes primarily from three species: *Strongylocentrotus purpuratus*, *Lytechinus variegatus*, and *Paracentrotus lividus*, all of which have the ancestral life history and diverged ∼35-50 million years ago (Figure 1A). Most regulatory interactions that have been experimentally tested in multiple species appear well conserved, as are expression domains of most of genes (McClay, 2011; Gildor and Ben-Tabou De-leon 2015; Israel et al. 2016; Massri et al. 2023).

We first assessed how well scRNA-seq is able to capture previously documented regulatory interactions in *Lv* by analyzing the distribution of regulator and target gene co-expression during development (Figures 7, S7, S8, S10; green dots and lines). In each case examined, co-expression corresponds to known developmental times and locations of specific regulatory interactions. For instance, *ets1* and *tbr* are expressed throughout the endomesoderm, but *ets1-sm32* and *tbr-foxB* are co-expressed exclusively within the skeletogenic cell lineage and only beginning at ∼12 hpf (Figure S9). Co-expression is readily detected for all gene pairs, despite the sparseness of scRNA-seq data. For most interactions, the majority of cells in the expected territory express multiple transcripts: dark green dots in the UMAPs indicate individual cells containing at least 2 unique molecular indices (UMIs) from each gene, while light green indicates just 1 UMI for one or both genes. Overall, co-expression plots are consistent with results from prior experimental studies and are sufficiently sensitive that absence of co-expression is biologically meaningful.

Based on this information, it is possible to make inferences about evolutionary conservation and change by examining co-expression of gene pairs among species (Figure S6). Comparisons of co-expression are shown in Figures 6, 7, S7, S8, S9, and S10 (*Lv* = green, *He* = orange). (1) *Conservation of an interaction*. Most gene pairs show co-expression in the same embryonic territories or cell types in both species. Co-expression in a completely distinct location from *Lv* was not observed in *He* for any of the gene pairs examined. The most straightforward interpretation of this pattern is that the ancestral regulatory interaction occurs during development in *He*. (2) *Temporal and/or spatial shift in a conserved interaction*. Although the location of co-expression was largely conserved in *He*, its timing and extent typically was not. Most shifts in timing involved a delay in the appearance of co-expression in *He* relative to *Lv*. Among many examples are *alx1-dri* (Figure S7) and *ets1-delta* (Figure S8). These cases are consistent with the general delay in specification and differentiation in *He* shown in Figures 3 and 4. While the timing of developmental gene expression can differ among sea urchin species with the ancestral life history, those shifts are typically smaller in magnitude and not biased in direction (Gildor and Ben-Tabou deLeon 2015; Israel et al. 2016; Massri et al. 2023). Clear examples of evolutionary differences in the extent of co-expression include all interactions specific to the gut and skeletogenic cells, both of which involve proportionally far fewer cells in *He* (Figures 7, S7, S8, S9, and S10). (3) *Loss of an interaction*. A minority of gene pairs that are co-expressed in the expected location in *Lv* show no or barely detectable co-expression in *He*. Examples include *alx1-foxB* (Figure S7) and *tbr-lasp1* (Figure S8). In these and other cases, lack of co-expression is not due to a technical issue with detection, as transcripts from both genes are detected elsewhere in the embryo (Figure S2). The most straightforward interpretation is that the specific regulatory interaction likely does not occur in *He*.

Inferred evolutionary changes in regulatory interactions in *He* are not randomly distributed across the developmental gene regulatory network, but instead concentrated around particular developmental processes. As discussed earlier, a very early patterning event in the ancestral dGRN is the establishment of cells that are both the founders of the skeletogenic cell lineage and the primary signaling center. In *Lv*, genes encoding the key transcriptional activators of the skeletogenic cell lineage, *alx1* and *ets1*, are first expressed at about the same time as genes encoding signaling ligands (*wnt1*, *wnt8*, and *delta*) (Massri et al. 2021). In contrast, expression of these genes occurs in two distinct phases and locations in *He*: an earlier phase in the archenteron involving genes that encode ligands (peaking at 9 hpf and greatly reduced by 12 hpf), and a later phase in the mesenchyme involving genes specific to skeletogenic cells (begins ∼16 hpf) (Figure 6B and S2). These results suggest that two key patterning events that are co-localized in the ancestral state have become independently regulated during the origin of the derived life history. This is remarkable, given the prior conservation of the ancestral state for over 230 million years (Thompson et al. 2015; Erkenbrack et al. 2018; Yamazaki et al. 2021).

The most obvious way a regulatory interaction could be lost during evolution is if the regulator is simply not expressed in the appropriate cell lineage or territory within the embryo. This is the case for two transcription factors, *tbr* and *foxB*. Both are expressed within the skeletogenic cell lineage of *Lv* (Saunders and McClay 2014), but the scRNA-seq data from *He* do not reveal any expression within these cells despite clear expression elsewhere in the embryo (Figure S2). Indeed, of the eleven genes known to encode transcription factors that activate expression within the skeletogenic cell lineage of species with the ancestral life history (Oliveri et al. 2008; Saunders and McClay 2014; Rafiq et al. 2012), *tbr* and *foxB* are only two that are not expressed in these cells in *He*. The absence of *tbr* expression may have limited impact on the expression of effector genes in skeletogenic cells, as Tbr appears to have far fewer targets than Alx1 and Ets1, the two primary activators of skeletogenic-specific transcription (Rafiq et al. 2012). The four known effector gene targets of Tbr are all expressed in skeletogenic cells of *He*, likely because they also receive input from other transcriptional activators, including Alx1 and Ets1 (Rafiq et al. 2012). *tbr* was previously proposed to be a more recent evolutionary addition to the skeletogenic cell GRN due to having fewer regulatory targets than Alx1 and Ets1 (Rafiq et al. 2012). The evolutionary loss of *tbr* expression within skeletogenic cells may have been possible for the same reason, coupled with the fact that nine other genes encoding transcription factors with roles in activating effector gene expression are also expressed within skeletogenic cells, thus providing some degree of regulatory redundancy.

### Evolution of morphology and life history

The evolution of massive maternal provisioning in *He* also precipitated changes in larval morphology and life history traits (Raff and Byrne 2006; Wray 2022). The most obvious are loss of feeding structures and a functional digestive tract (Williams and Anderson 1975), which are no longer needed with a richly provisioned egg (Hoegh-Guldberg and Emlet 1997; Byrne et al. 1999; Davidson et al. 2019). Another set of changes was likely driven by selection to reduce larval mortality by shortening pre-metamorphic development, including earlier left-right symmetry breaking, differentiation of coeloms, and formation of the adult imaginal rudiment (Williams and Anderson 1975; Wray and Raff 1989; Henry et al. 1991; Koop et al. 2017).

The scRNA-seq data reflect both sets of changes in the proportions of cell types in the early larva (Figure 1D). In *He*, far fewer cells are allocated to endoderm, which is non-functional until after metamorphosis, and to skeletogenic cells, which produce a vestigial larval skeleton (Williams and Anderson 1975; Emlet 1995). Conversely, more cells are allocated to coeloms and ectoderm, both of which contribute substantially to accelerated development of the post-metamorphic juvenile (Williams and Anderson 1975; Wray and Raff 1989; Koop et al. 2017).

In sea urchins with the ancestral life history involving feeding larvae, four territories of ectodermal cells are evident from anatomy and gene expression: a ciliated band used for feeding and locomotion, an anterior neurogenic domain, and generalized ectoderm with distinct oral and aboral domains. These territories are recovered as separate clusters with scRNA-seq in *Lv* (Figures 1B and S3). Previous studies examining ectodermal gene expression in *He* found no evidence of conserved oral and aboral territories, and suggested instead that the ancestral ectodermal domains are reorganized (Haag and Raff 1998; Love and Raff 2006; Koop et al. 2017). The scRNA-seq data reveal a well-defined anterior neurogenic domain in *He* (Figures S2 and S3). However, the other ancestral ectodermal domains are more difficult to recognize in the *He* larva. Markers of oral and aboral ectoderm the ancestral state are not consistently co-localized in *He* (Figure S3). The ciliated band, which is used for feeding, has been lost in *He* (Williams and Anderson, 1975). The only regions of dense cilia in the *He* larva likely correspond instead to the epaulettes of late larvae in the ancestral state, which are used exclusively for swimming (Emlet 1995). The other derived trait in *He* that likely contributes to changes in expression of regulatory genes within the ectoderm is the greatly accelerated development of the imaginal adult rudiment (Williams and Anderson 1975; Wray and Raff 1989; Emlet 1995; Koop et al. 2017). Vestibular ectoderm is a distinct gene expression territory within the ectoderm of *He* by 24 hpf (Koop et al. 2017). Comparative analysis of gene expression in the epaulettes and vestibule will require extending the *Lv* scRNA-seq time course to late larval stages, as these structures have not yet developed in the early larva.

In summary, the scRNA-seq data presented here reveal numerous features of development in *He* that are likely conserved and others that are likely modified since its divergence from other sea urchins that share the ancestral life history. While scRNA-seq data alone do not provide direct evidence about molecular mechanisms, they can produce detailed information about specific developmental processes that are not evident from bulk RNA-seq and would otherwise require gene-by-gene expression analyses. Here we report a close correlation between evolutionary changes in the timing and location of regulatory genes and evolutionary changes in larval morphology and life history. Many specific regulatory interactions that are widely conserved among sea urchins with the ancestral life history appear to be conserved but delayed in *He*, while a small number may have been lost entirely. These results provide specific predictions that can be tested efficiently using perturbation experiments, greatly facilitating the daunting challenge of understanding which connections within developmental gene regulatory networks are conserved, altered, or lost during the course of evolution.

## ACKNOWLEDGEMENTS

We thank the staff of the Sydney Institute for Marine Sciences for skilled support and the staff of Duke’s Sequencing and Genomic Technologies core facility for expert advice and assistance. Echinobase was critical to the success of this project. Matthew Clements, Phillip Davidson, Hannah Devens, Allison Edgar, Brennan McDonald, Carl Manner, Esther Miranda, Zach Pracher, Paulina Selvakumaraswamy, Jane Swart, and Emma Wallace provided invaluable advice. This research was financially supported by NIH Training Grant HD040372 and NSF Graduate Research Fellowship DGE-1644868 to AJM; NIH grant HD014483 to DRM; and NSF grant IOS 1929934 to GAW.

## METHODS

### Methods

#### Spawning and embryo culture

Adult *H. erythrogramma* were collected under permit near Sydney, Australia during October and November. Crosses were initially established for the purpose of optimizing dissociation protocols; subsequently, a single cross was used to source samples for this study. Adults were spawned by injecting 0.5 ml 0.5 M KCl intracoelomically. Unfertilized eggs were allowed to float and washed 3X in filtered natural sea water (FNSW). Eggs were fertilized with sperm in FNSW containing 0.02 g PABA / 100 ml. Zygotes were washed an additional 3X in FNSW to remove residual sperm and PABA and embryos cultured at 23 °C in FNSW. At each time point, embryos were visually verified to be morphologically similar prior to dissociation. Throughout, methods closely matched our previous scRNA-seq analysis of *L. variegatus* (Massri et al. 2021), including only slightly species-optimized dissociation protocols, same rearing temperature, time-matched samples, and same versions of 10X library kits and Illumina sequencing chemistry.

#### Time points sampled

Embryos / larvae were sampled at seven time points: 6, 9, 12, 16, 20, 24, and 30 hpf (late cleavage through early larva). Time points were chosen to align with Massri et al. 2021, with two additional considerations. First, due to the large egg size of *He* (∼430 µm diameter), blastomeres exceed the diameter of the microfluidics on the 10X platform until the 512-cell stage (6 hpf), which became our first time point. Second, prior studies suggested that activation of the zygotic genome in *He* is somewhat delayed relative to *Lv* (Wang et al. 2020; Davidson et al. 2022); thus, we collected one additional time point (30 hpf) beyond the last sampled in *Lv* (24 hpf). Comparative analyses, drew on published data from Massri et al. 2021 for *Lv* and from the present study for *He*.

#### Cell dissociation and fixation

At each time point, the culture was sub-sampled and embryos washed two times in Calcium-Free Artificial SeaWater (CFASW). ∼3ml embryos in CFASW were added to 7ml of dissociation buffer (1.0M glycine and 0.25 mM EDTA, pH 8.0 with HCl) at 4 °C, and then placed on a rocker for 4 minutes. Following incubation, samples were triturated 10-15X, then 10ml of ice-cold methanol was added, and incubated for 4 additional minutes on a rocker. Following incubation, samples were triturated 10-15X additionally, and visually inspected under a microscope for a homogenous single cell suspension. To fix cells, 40ml of ice-cold methanol was added to a final concentration of 80%. Samples were then placed on a rocker for one hour prior to storage at -20°C.

#### Library preparation and sequencing

Fixed cells were washed once in methanol, then rehydrated by washing in 3X sodium citrate buffer. Cell concentrations were determined using a hemacytometer. Seven libraries were prepared using the 10X Genomics 3 ’v3 gene expression kit and the 10X Chromium platform to encapsulate single cells within droplets. Library quality was verified using an Agilent 2100 Bioanalyzer. Libraries were titered and pooled at Duke University’s Sequencing and Genomic Technologies Core Facility, then sequenced in one S1 flow cell on an Illumina NovaSeq 6000 with 28 x 8 x 91 bp.

### Computational Analysis

#### Initial processing and production of raw csv count files

Following sequencing, Cellranger 3.1.0 was used to convert Illumina-generated BCL files to fastq files using the Cellranger “mkfastq” command. The “mkref” command was then applied to index the *H. erythrogramma* 1.0 Genome (Davidson et al. 2022a). The “count” command was used to demultiplex and quantify reads mapping to the reference *He* genome. The “mat2csv” command was used to generate CSV RNA count matrix files for each time point for downstream analysis.

#### Data filtering and normalization

All 19 *Lv* and 7 *He* CSV RNA count matrix files were uploaded to R, and a merged Seurat object (Hao et al. 2022) was generated for each species. The *Lv* Seurat object was filtered to remove lower quality cells with nFeature_RNA > 200, nFeature_RNA < 7000, and nCount_RNA < 10000. In total, 50,638 *Lv* cells remained. The *He* Seurat object was filtered with nFeature_RNA > 200, nFeature_RNA < 4000, and nCount_RNA < 10000. In total, 23,156 *He* cells remained. SCTransform was then independently applied to the *Lv* merged filtered object and the *He* merged filtered object to perform normalization, and regression of ribosomal and cell cycle related genes using the command: *vars.to.regress = c(“percent.Rb”, ”cell.cycle”)*. These metacolumns were added with the following commands: *PercentageFeatureSet(merged, pattern = “\\b\\w*Rp[sl]\\w*\\b”, col.name = “percent.Rb”) and PercentageFeatureSet(merged, pattern = “\\b\\w*C[d|y]c\\w*\\b”, col.name = “cell.cycle”)*.

#### Dimensionality reduction, visualization, and clustering

We next independently performed Principal Component Analysis on the SCTransformed *Lv* Seurat object and *He* Seurat object, and found the nearest neighbors and clusters (Hao et al. 2022). UMAP was then applied to each species to visualize the multidimensional scRNA-seq in a two-dimensional space. Each species clusters were annotated using co-expression of dGRN genes, and published *in situ* hybridization patterns as markers. Echinobase (Arshinoff et al. 2021) was used to identify gene function. See Massri et al. 2021 for a list of marker genes and supporting literature.

#### Multi-species integrated analysis

Orthologroups were identified using OrthoFinder v 2.5.4 (Ems and Kelly 2019) and used to generate a list of 1:1 orthologues in *Lv* and *He*. In total, 7,349 genes were identified as 1:1 orthologues and expressed in the combined data set of 72,445 cells. The standard Seurat/SCTransform pipeline was performed, and then integrated by species using the Canonical Correlation Analysis (CCA) workflow (Butler et al. 2018) using *Lv* as the reference.

#### Waddington-OT developmental trajectories

To infer developmental trajectories in *He*, we used Waddington-OT (Schiebinger et. 2019). To execute we used the SCTransform normalized expression matrix obtained after running Seurat, a table of cell barcodes with cell-type annotations, and a growth rates table that was estimated from expected changes in lineage proportions over time using the model implemented in Waddington-OT. To estimate cell division rates, we used the best estimate of the expected number of cells at key developmental time points. We assumed that cell divisions were uniform between estimates of expected cell numbers. Next we recalculated transport maps using the modeled cell division rates, optimization parameters ε=0.05, λ1=1, and λ2=50, and 20 iterations of growth rate learning. We used the transport map model throughout our analysis that included triangle plots and lineage trees.

#### Waddington-OT time alignment

To estimate timing differences between the *Lv* and *He* datasets, we used optimal transport combined with the gene orthology tables. First, we used the previously calculated transport maps for both datasets to obtain fate probabilities for the cells at each time point. Fate probabilities were computed relative to cell types found in the last time point of their respective dataset. Next, we restricted the normalized counts for *Lv* and *He* to known gene orthologues using the previously generated gene orthologue table. Then, for each cell type, a time point by time point matrix of earth mover distances between the two datasets was computed. In the calculations for each cell type and pair of times, cells were weighted by their fate probabilities to the cell type in question. Finally, for each *Lv* time point, the *He* time point corresponding to the minimum earth mover distance to it was found. These pairs were found for each cell type. We then take these pairs of time points to be the optimal developmental time alignments for the cell type.

#### Waddington-OT triangle plots

To construct triangle plots, we used transport maps calculated before to compute fate probabilities with respect to the last common time point in our dataset (24 hpf) and visualized them by computing the barycentric coordinates of cell fates between two different cell types and at a threshold of 0.7.

#### Developmental lineage trees

To infer cell lineage trees, we used our modeled transport maps to find connections between cell clusters by calculating the fraction of descendants that end up in cluster/cell-type *j* at time *ti+1* from cluster/cell-type *i*. The minimum number of cells for a cluster to be represented set to 10 and the minimal edge weight cutoff was set to 0.15. Once the unwanted edges were removed, the data was written in a format that is usable by d3.js.

#### Co-expression analyses

The number of cells expressing any two specified genes was tallied from the count tables and normalized by the total number of cells for each sample using a custom Python script, and used to generate time courses for both species. To visualize co-expression in UMAP space, a custom R script was used.

## SUPPLEMENTARY FIGURES AND TABLES

**Figure S1. Metrics of scRNA-seq transcriptomes over development in *He*.** Violin plots showing the frequency distribution of four informative metrics. **A.** Distinct genes/cell from which transcripts were detected (nFeature_RNA in Seurat). The modest decline over development likely reflects the transition of a more restricted transcriptome during differentiation. **B.** Distinct UMIs recovered per cell (nCount_RNA in Seurat). **C.** Percentage of transcripts mapping to genes encoding ribosomal proteins. The increase over development reflects the maternal-to-zygotic transition in protein expression. **D.** Percentage of transcripts mapping to genes encoding cell cycle control proteins. The slight decrease in the last two stages reflects the slowing of cell division as differentiation commences in the early larva.

**Figure S2. Expression of marker genes in *He.*** Density plots showing the distribution in UMAP space of cells expressing marker genes for specific embryonic territories and larval cell types. See Massri et al. 2021 for supporting literature.

**Figure S3. Expression of marker genes for ectodermal territories in *Lv* and *He*.** Density plots showing the distribution in UMAP space of cells expressing marker genes for specific embryonic territories and larval cell types. See Massri et al. 2021 for supporting literature.

**Figure S4. Overall similarity of transcriptomes in *Lv* and *He* over development.** Heatmap showing overall similarity of expression of 1:1 orthologues among all developmental stages examined. Similarity measured using CIDER (Hu et al. 2021).

**Figure S5. Evolutionary changes in timing of differentiation, additional examples.**

Optimal transport was used to predict the likely fate for each cell at five stages, based on transcriptomes at 24 hpf (see Methods). The organization of this figure parallels that of main text Figure 4. Triangle plots show transcriptomes predictive of two specific cell types. **A.** Coelom (red) and endoderm (yellow). **B.** Blastocoelar cells (purple) and endoderm (yellow). **C.** Blastocoelar cells (purple) and pigment cells (pink). Any other cell fate is indicated in dark gray; cells with undifferentiated transcriptomes are shown in blue. See text for interpretation.

**Figure S6. Logic underlying inference of distinct types of evolution change in regulatory interactions.** On the extreme left, hypothetical data from *Lv* illustrating an experimentally validated regulatory interaction: transcriptional activator, gene A (blue) and its target, gene B (purple), are co-expressed in the same cells. Four possible scenarios in *He* for the same pair of genes are shown to the right, with interpretations regarding the regulatory interaction shown below. Scenario 1 is consistent with conservation. Scenario 2 is consistent with conservation, but with a delay in the timing of the interaction. Scenario 3 implies that the interaction no longer occurs and that some other transcription factor must activate expression of gene B. Scenario 4 is consistent with conservation, but additionally implies that some other transcription factor must operate earlier to initiate expression of gene B.

**Figure S7. Co-expression of *alx1* and several of its interactors within the skeletogenic dGRN. See caption to Figure 7**.

**Figure S8. Co-expression of additional regulators and targets within the skeletogenic dGRN. See caption to Figure 7**.

**Figure S9. Comparison of regulator expression and co-expression with a specific target gene.** Five validated transcription factor → target gene interactions are illustrated. Left panels show density plots of expression for the regulator while right panels show cells expressing both the regulator and its target. Note that co-expression domains are often a subset of the cells expressing the regulator.

**Figure S10. Co-expression of several regulators and targets in non-skeletogenic territories of the dGRN. See caption to Figure 7**.

Table S1. Cell counts over development in *Lv*.

Table S2. Cell counts over development in *He*.

## Notes

### Competing Interest Statement

The authors have declared no competing interest.

