## Supplementary figures and images for "Single-cell transcriptomics reveals evolutionary reconfiguration of embryonic cell fate specification in the sea urchin *Heliocidaris erythrogramma*"

### Supplemental Figure 1

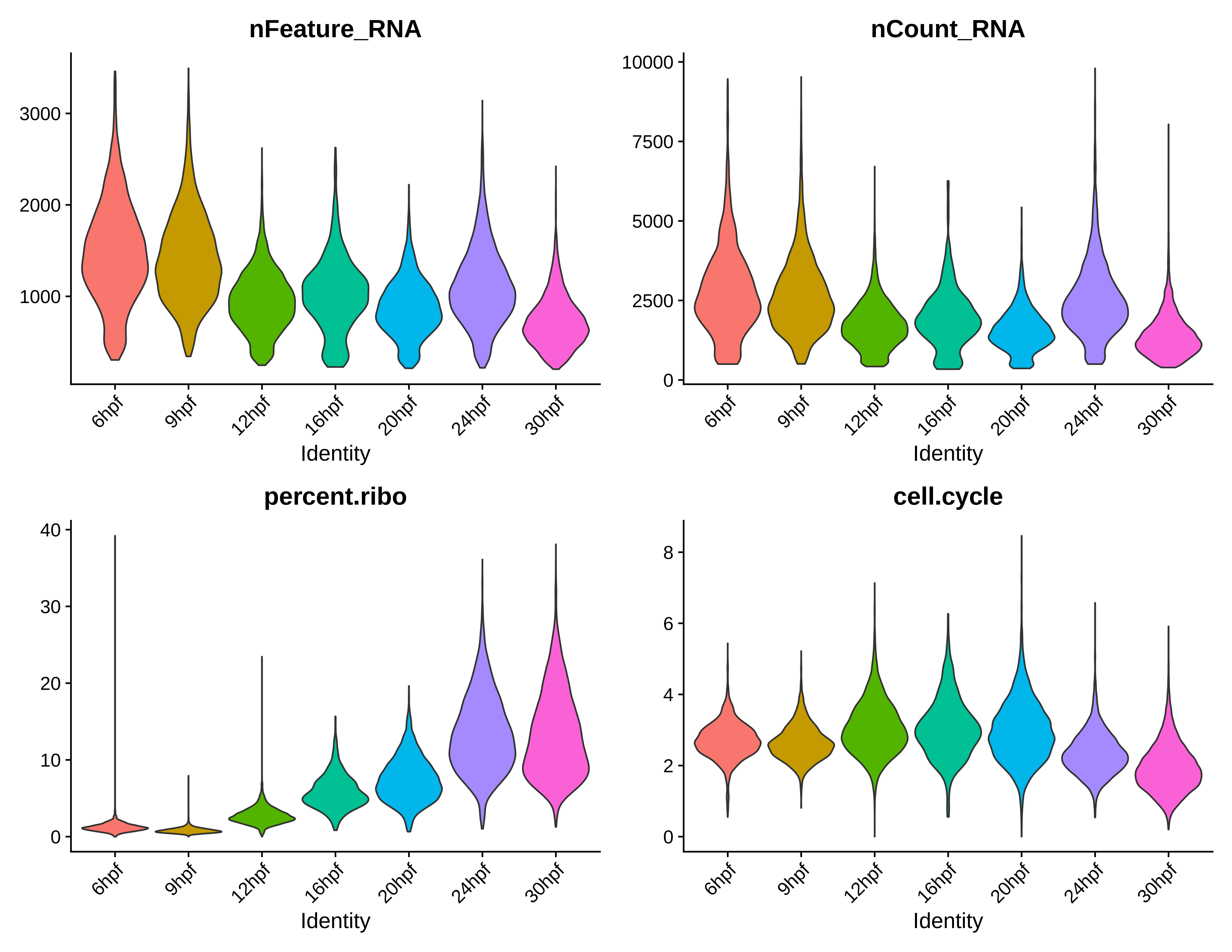

### Supplemental Figure 2

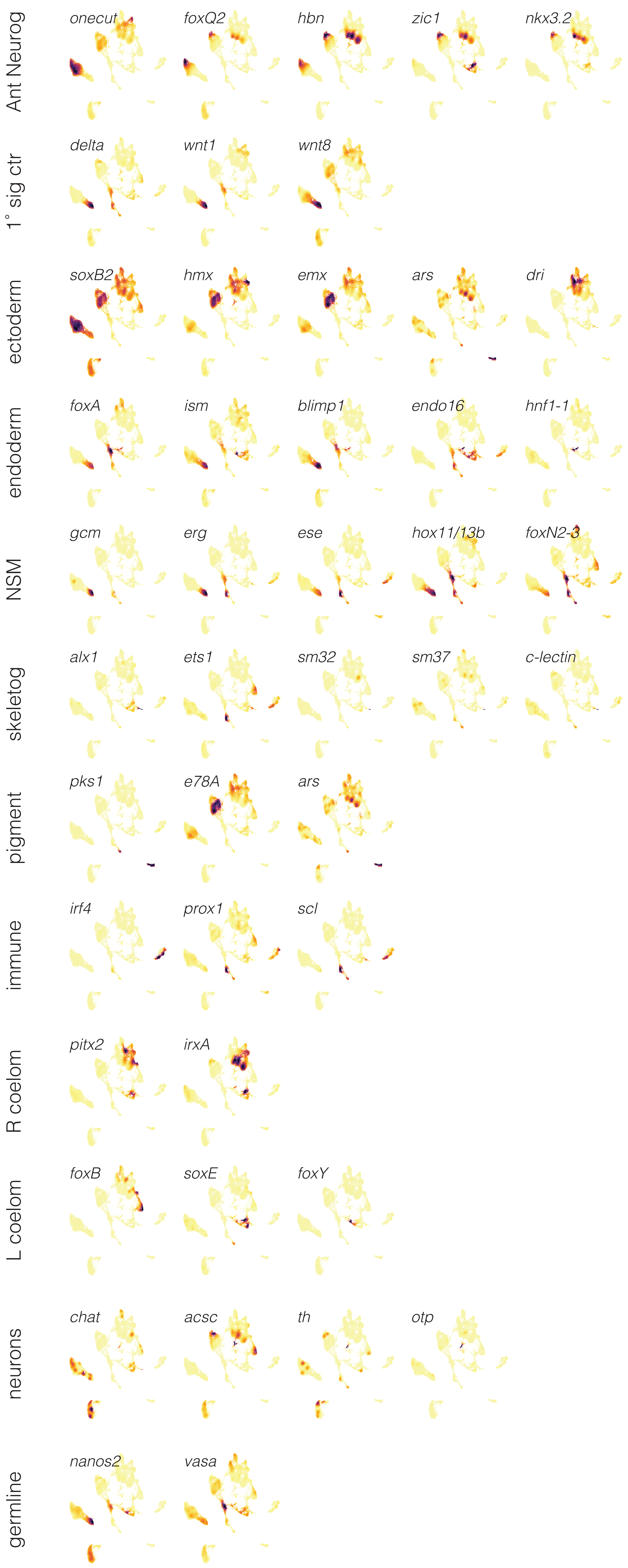

### Supplemental Figure 3

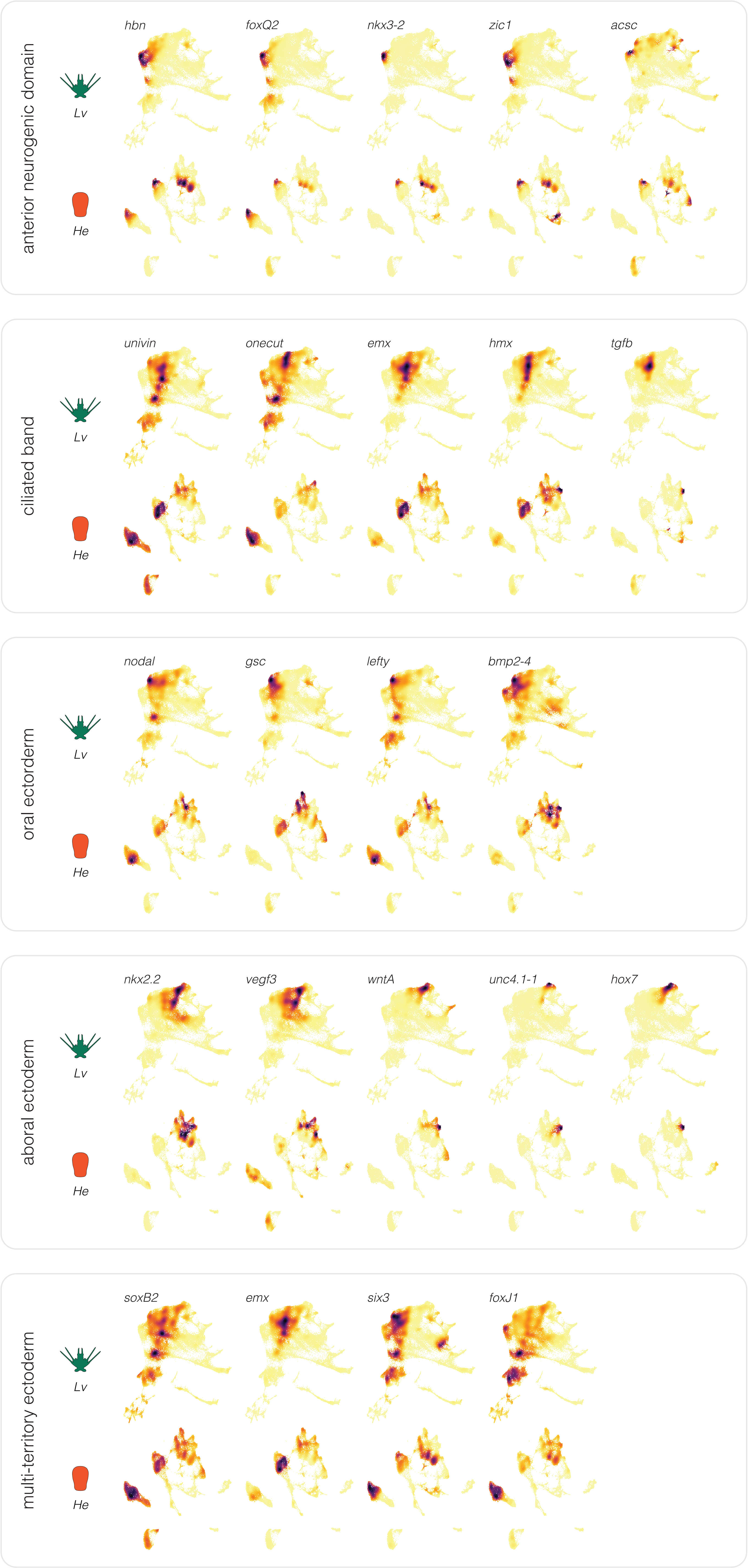

### Supplemental Figure 4

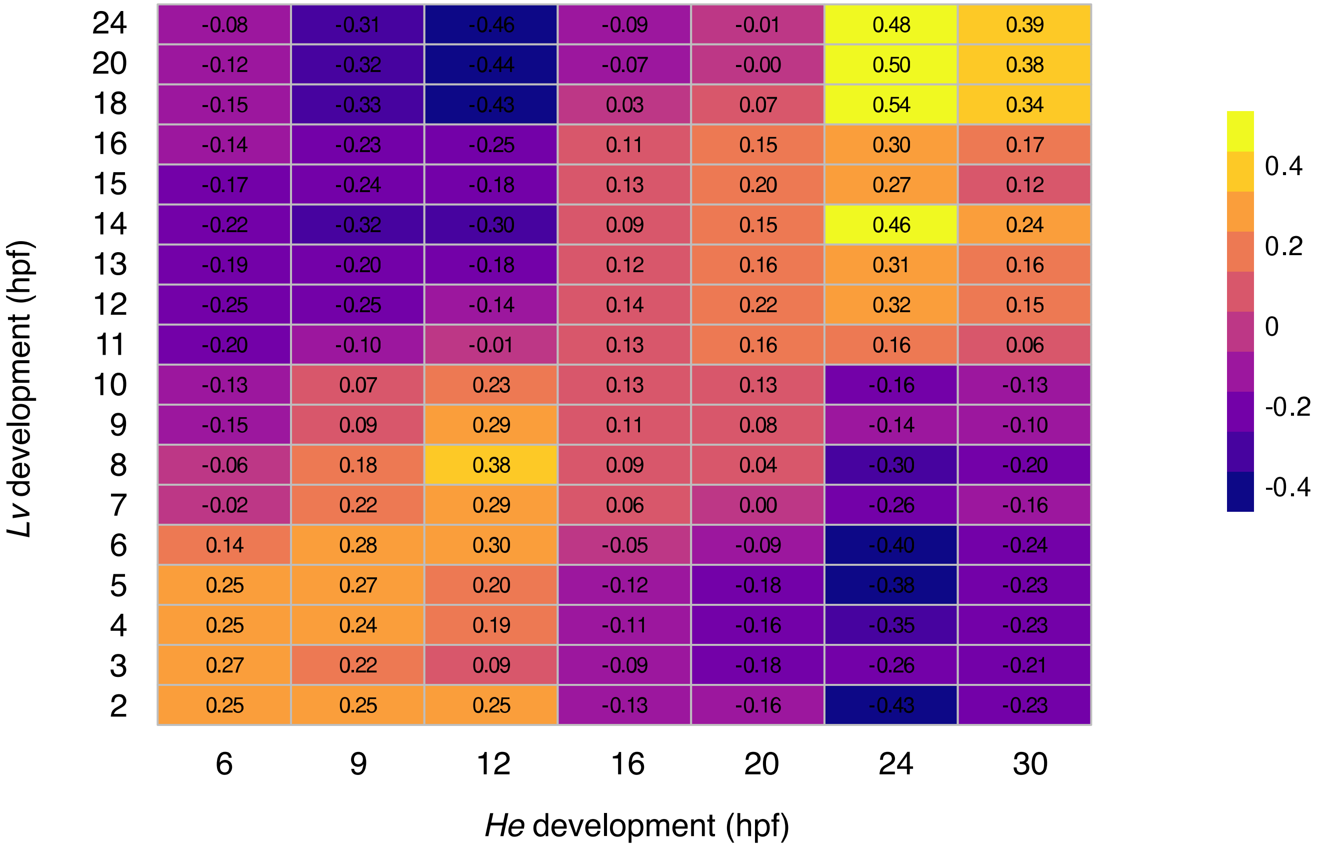

### Supplemental Figure 5

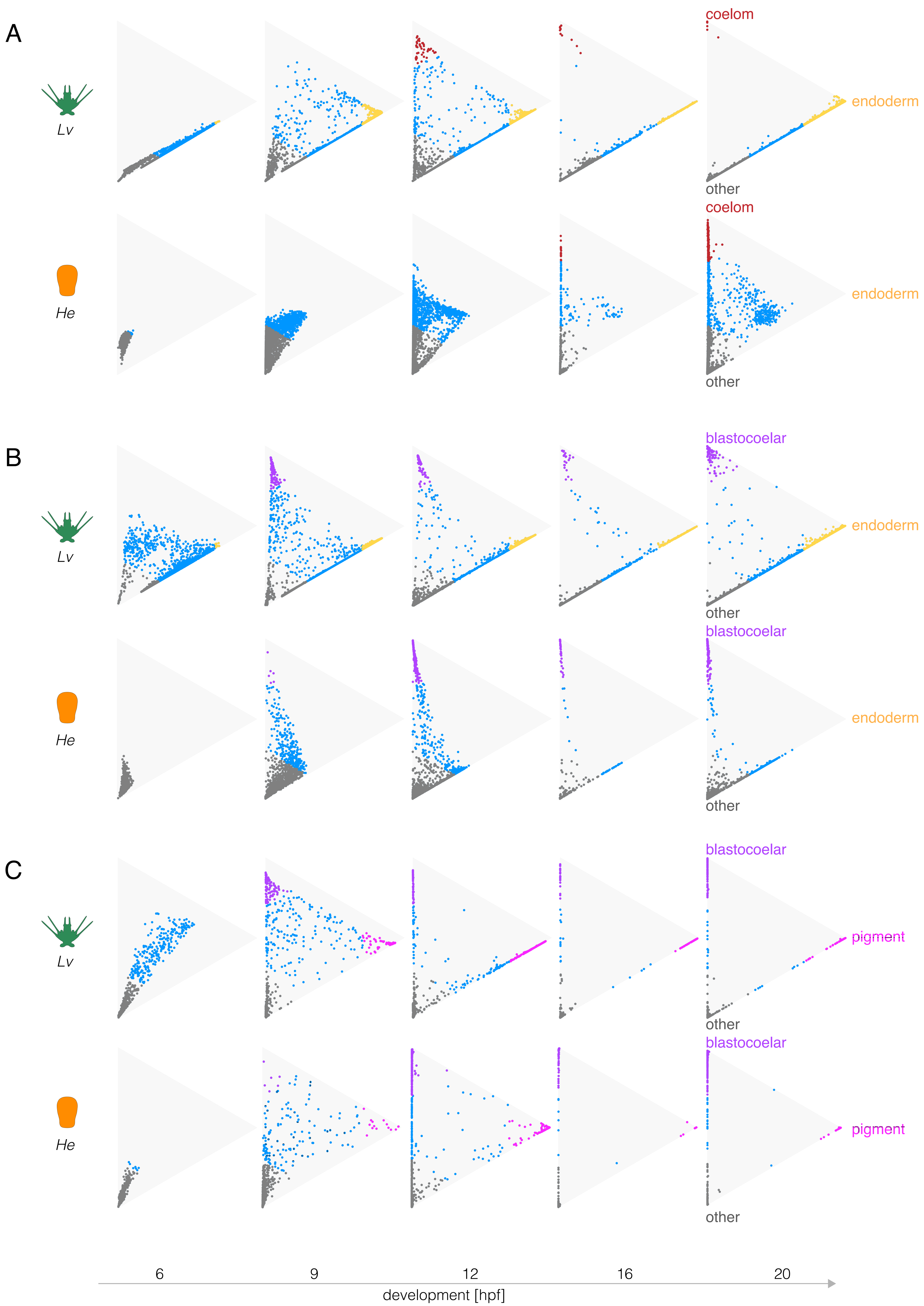

### Supplemental Figure 6

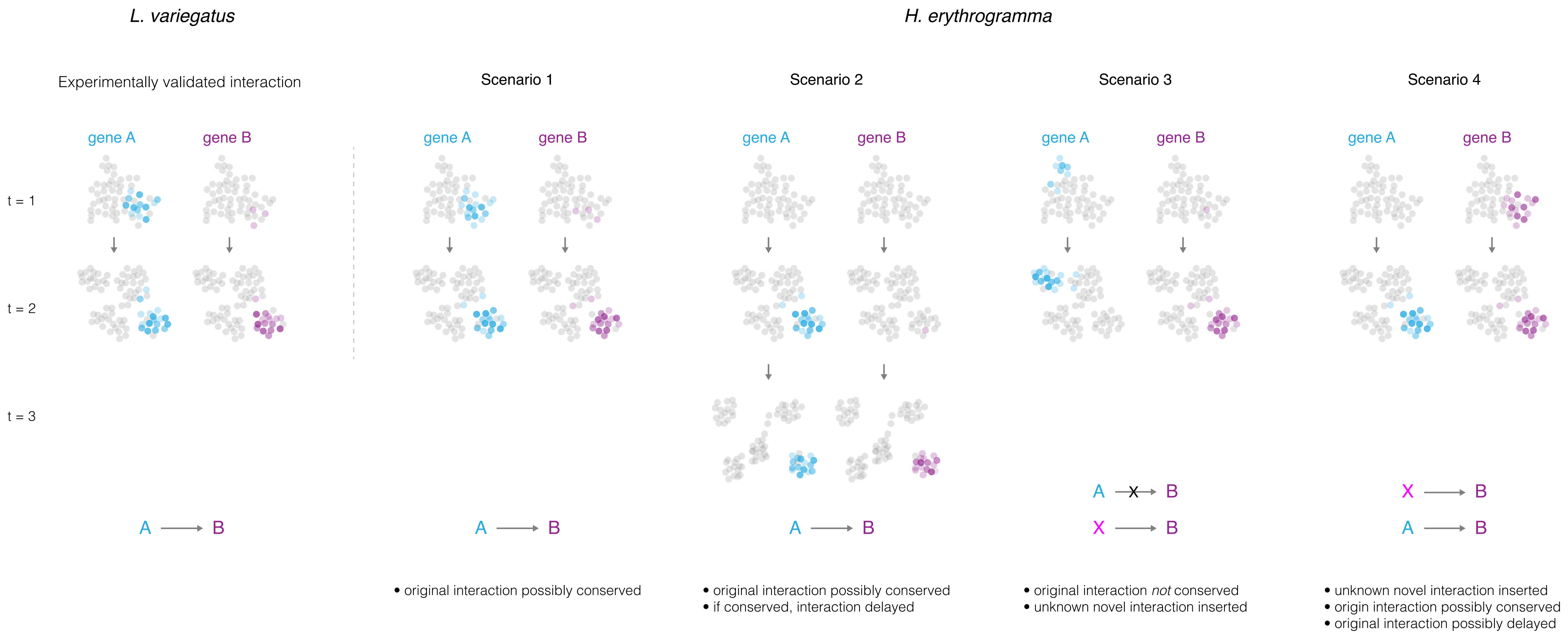

### Supplemental Figure 7

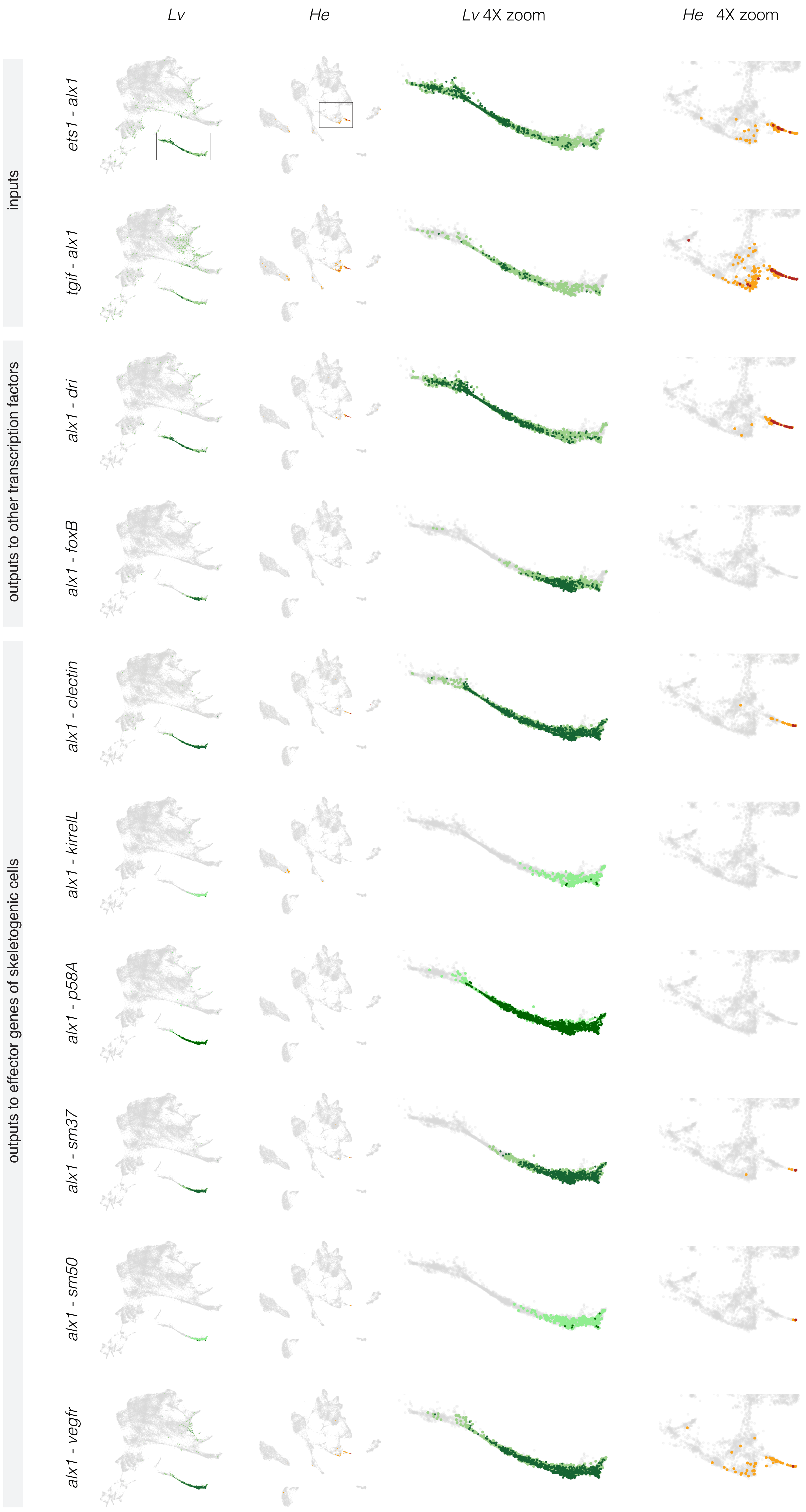

### Supplemental Figure 8

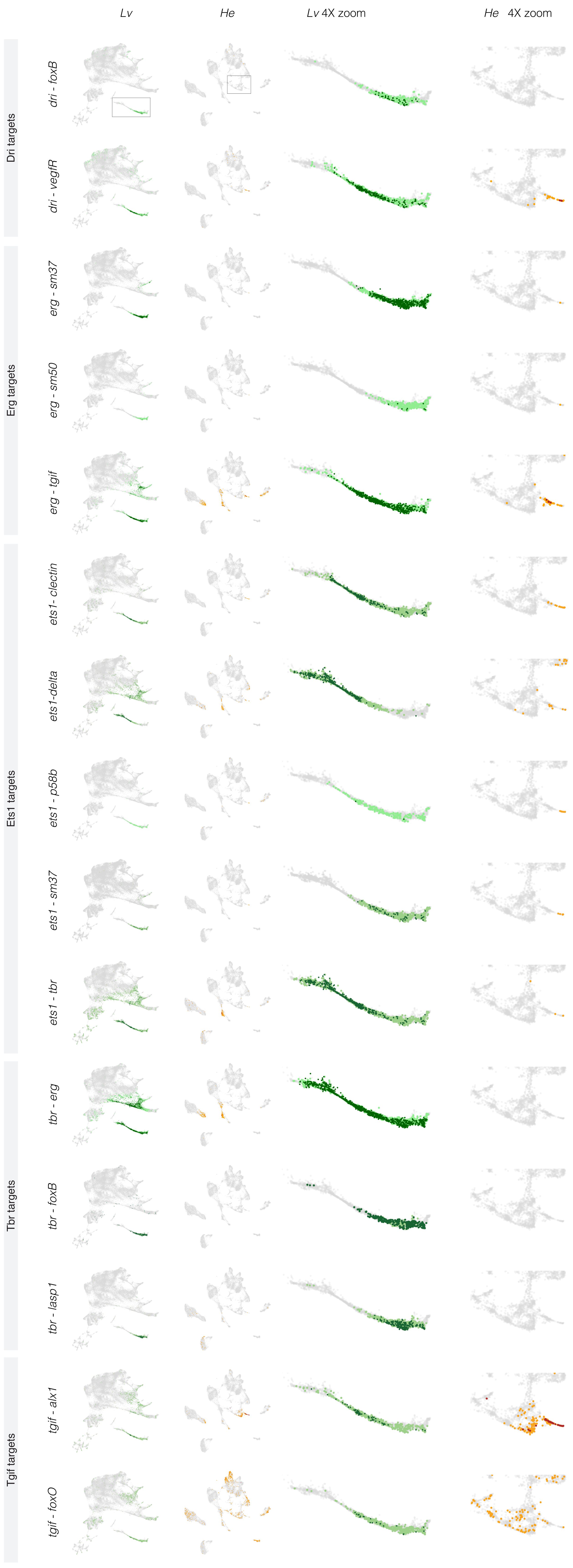

### Supplemental Figure 9

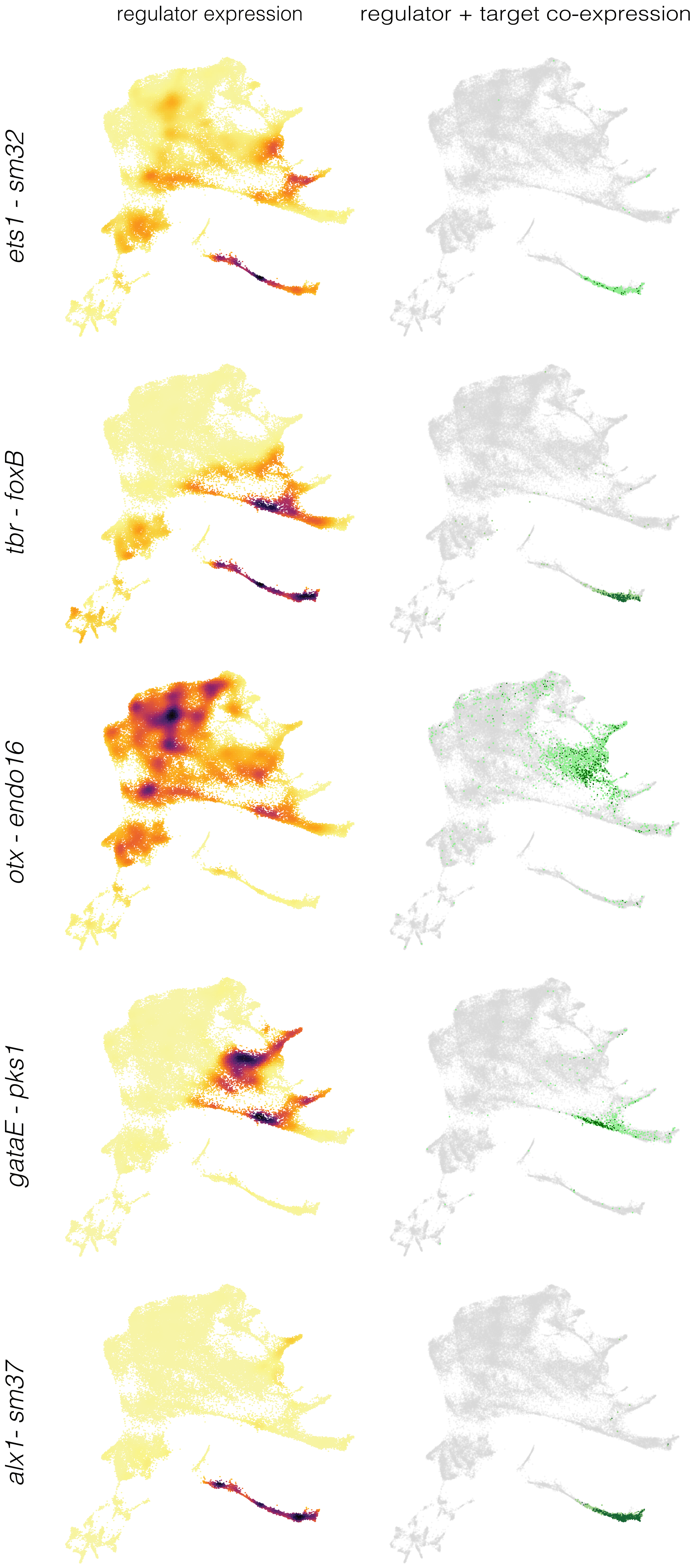

### Supplemental Figure 10

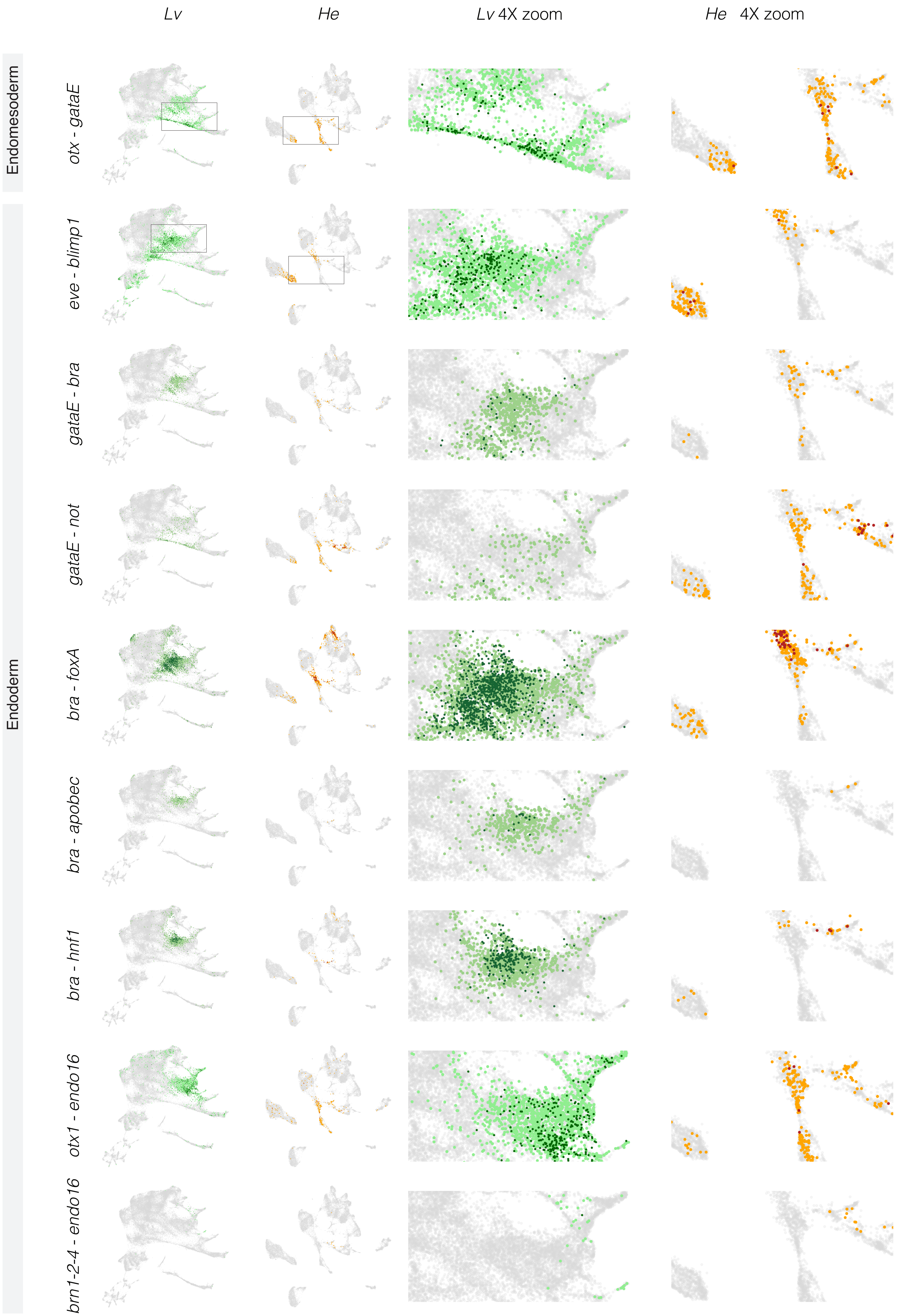
